# Genetic structure, function and evolution of capsule biosynthesis loci in *Vibrio parahaemolyticus*

**DOI:** 10.1101/2020.02.25.964247

**Authors:** Shengzhe Bian, Zeng Wenhong, Qiwen Li, Yinghui Li, Nai-Kei Wong, Min Jiang, Le Zuo, Qinghua Hu, Liqiang Li

## Abstract

Capsule-forming extracellular polysaccharides are crucial to bacterial host colonization, invasion, immune evasion and ultimately pathogenicity. Due to warming ocean waters and human encroachment of coastal ecosystems, *Vibrio parahaemolyticus* has emerged as a globally important food-borne enteropathogen implicated in acute gastroenteritis, wound infections, and septic shock. Conventionally, the antigenic properties of lipopolysaccharide (LPS, O antigen) and capsular polysaccharide (CPS, K antigen) have provided a basis for serotyping *V. parahaemolyticus*, while disclosure of genetic elements encoding 13 O-serogroups have allowed molecular serotyping methods to be developed. However, the genetic structure of CPS loci for 71 K-serogroups has remained unidentified, limiting progress in understanding its roles in *V. parahaemolyticus* pathophysiology. In this study, we identified and characterized the genetic structure and their evolutionary relationship of CPS loci of 40 K-serogroups through whole genome sequencing of 443 *V. parahaemolyticus* strains. We found a distinct pattern of CPS gene cluster across different K-serogroups, and expanded its new right-border by identifying *glpX* as a key gene conserved across all serotypes. A total of 217 genes involved in CPS biosynthesis were annotated. Functional contents and genetic structure of the 40 K-serogroups were analyzed. Based on inferences from species trees and gene trees, we proposed an evolution model of the CPS gene clusters of 40 K-serogroups. Horizontal gene transfer by recombination from other *Vibrio* species, gene duplication and nonsense mutations are likely to play instrumental roles in the evolution of CPS in *V. parahaemolyticus.* It is the first time, to the best of our knowledge, that a large-scale of CPS gene clusters of different K-serogroups in *V. parahaemolyticus* have been identified and characterized in evolutionary contexts. This work should help advance understanding on the variation of CPS in *V. parahaemolyticus*, and provide a framework for developing diagnostically relevant serotyping methods.

**Author summary:** Due to warming ocean waters and human encroachment of coastal ecosystems, *Vibrio parahaemolyticus* has emerged as a globally important food-borne enteropathogen. However, the genetic structure of CPS loci for 71 K-serogroups *V. parahaemolyticus* have remained unidentified, limiting progress in understanding its roles in *V. parahaemolyticus* pathophysiology. In this study, we identified and characterized the genetic structure of CPS loci of 40 K-serogroups through whole genome sequencing of 443 *V. parahaemolyticus* strains. We expanded and identified its new right-border by identifying *glpX* as a key gene conserved across all serotypes. We proposed an evolution model of the CPS gene clusters of 40 K-serogroups. We also found horizontal gene transfer by recombination from other *Vibrio* species, gene duplication and nonsense mutations are likely to play instrumental roles in the evolution of CPS in *V. parahaemolyticus.* It is the first time, to the best of our knowledge, that a large-scale of CPS loci of different K-serogroups in *V. parahaemolyticus* have been identified and characterized in evolutionary contexts. This work should help advance understanding on the variation of CPS in *V. parahaemolyticus*, and provide a framework for developing diagnostically relevant serotyping methods.

## Introduction

*Vibrio parahaemolyticus*, a Gram-negative halophilic bacterium, is taxonomically a notable member of the genus *Vibrio* within the family *Vibrionaceae*. It prevails in estuarine, marine and coastal areas [1–4], and is typically isolated in a free-swimming state. Significant motility is conferred by a single polar flagellum in *V. parahaemolyticus*, which is capable of sensing both biotic and abiotic surfaces including zooplankton, fish, and shell-fish, via impeded rotation [5]. Due to climate change and anthropogenic degradation of coastal environments, *V. parahaemolyticus* is gaining notoriety as an enteropathogen in humans worldwide, where people depend on seafoods as a major nutritional source. Clinically, it can cause three major syndromes, namely: gastroenteritis, wound infections, and septicemia [6]. In general, thermostable direct hemolysin (TDH) and TDH-related hemolysin (TRH) are two major virulence factors of *V. parahaemolyticus* implicated in its pathogenicity [7].

In host-pathogen interactions, bacteria leverage several extracellular polysaccharides to colonize hosts and cause disease. Gram-negative bacteria produce lipopolysaccharide (LPS), which is an integral component of the outer leaflet of bacterial outer membrane consisting of three structural moieties: lipid A, core oligosaccharide (core), and O-specific polysaccharide or O antigen (OAg) [8, 9]. An additional capsular polysaccharide (CPS) (K antigen) may also be expressed, which is considered a significant virulence factor as it can increase bacterial survival upon phagocytosis by eukaryotic cells [10, 11] and blunt efficacy of antibiotics [12, 13]. Importantly, capsule is also advantageous for bacterial persistence and adaptation to harsh environments through protection from physical and chemical stresses, without sacrificing efficiency in the transfer of genetic materials between cells [14].

As a Gram-negative bacterium, *V. parahaemolyticus* produces a number of different somatic (O) and capsular (K) antigens, which have been exploited as a primary basis of strain classification [15]. Being a pathogenic bacterium of multi-serotypes, *V. parahaemolyticus* can be classified into 13 O serotypes and 71 K serotypes [16]. Progress on the variation and dissemination of those serotypes have gone through different stages in its research history. *V. parahaemolyticus* was first discovered by Tsunesaburo Fujino in 1950 as a causative agent of food borne disease following a large outbreak in Japan, which recorded 272 patient cases with 20 deaths after consumption of *shirasu* [17]. Before 1996, no particular serotypes of *V. parahaemolyticus* were associated with outbreaks. However, in the same year, a major outbreak arose in Kolkata, India, later known as the first pandemic, which was caused by strains with increased virulence. More than half of the patient isolates were serotype O3:K6 [15]. Subsequently, the pandemic serotypes disseminate widely and rapidly. Within a few months, pandemic O3:K6 strains were identified in neighboring Vietnam, Indonesia, Bangladesh, Laos, Japan, Korea and Thailand [15]. They have consistently been detected globally (including Africa, Europe, North America and Latin America) for years afterward [15, 18, 19]. A variety of serotype variants have emerged such as O4:K68, O1:K25 and O1:KUT. However, they have identical molecular characteristics similar to the pandemic O3:K6 and have thus been collectively referred to as sero-variant of pandemic O3:K6 strains [20]. Until 2016, a total of 49 pandemic serotypes (including 30 K-serogroups without untypeable) isolates from 22 countries across four continents (Asia, Europe, America, and Africa) were identified. All of these serotypes were detected in clinical isolates. Notably, due to its large geographical span and population, China has the most abundant pandemic serotypes among the 22 affected countries, which reach up to 26 serotypes and 12 K serogroups without untypeable [21].

Traditionally, serotyping assays use commercially available antisera to identify *V. parahaemolyticus* strains but this approach is limited by high costs, complicated procedure, cross-reactivity and even subjective interpretation [22]. In contrast, molecular methods targeting serotype-specific genes can circumvent these shortages with proven superiority in specificity and sensitivity in the identification of bacterial serotypes [23]. For instance, molecular methods for 13 O-serogroups detection and identification for *V. parahaemolyticus* have been developed, based on specific genes of O-serogroup genetic determinants (OGDs) [24]. However, the CPS gene cluster that determines K-serogroup has not been reliably identified to this date. In fact, there is a paucity of knowledge on the location of CPS gene cluster in the *V. parahaemolyticus* genome, which was controversial up until 2010. For the pandemic O3:K6 strains, Guvener *et al.* proposed a locus VPA1403-VPA1412 on chromosome II for capsular polysaccharide biosynthesis without experimental verification of its functional correlation with K-antigen [25]. Subsequently, through homologous alignment with O-side chain and core OS loci in *Vibrio cholera* and comparison of restriction fragment polymorphisms in different serotypes, Okura *et al*. suggested a different locus determinate for K-antigen, specifically, around VP0214-VP0238 (*gmhD* – *rjg*) on chromosome I, though experimental evidence was not furnished [26]. In 2010, Yuansha Chen *et al*. investigated these putative K-antigen genetic determinants in a pandemic O3:K6 isolate and confirmed by gene deletion that VP0214-VP0238 (*gmhD* – *rjg*) on chromosome I determines K-antigen specifically but not O-antigen [27]. Although location of K-antigenic determinant has been proven in pandemic O3:K6, the genetic structure and function of the loci encoding other K-antigens remain obscure. Previous studies have found that most prevalent clones of *V. parahaemolyticus* stemmed from pandemic O3:K6 via serotype conversion [15, 20, 28–30], while pandemic *V. parahaemolyticus* serotypes have become more and more diverse since 1996 [15]. In order to advance a better understanding of *V. parahaemolyticus* pandemics, it is imperative to clarify the evolutionary relationship and divergence mechanism of CPS loci across different *V. parahaemolyticus* K-serogroups.

In this study, we have identified CPS loci of 40 K-serogroups from 64 serotypes, which include 24 pandemic K-serogroups (covering 86% pandemic K-serogroups in the world and 92% pandemic K-serogroups in China) and 16 non-pandemic K-serogroups. Their genetic structure, function and evolutionary relationship of these 40 K-serogroups CPS loci were investigated. This work provides a framework for analyzing frequent mutations in *V. parahaemolyticus* K-antigens and developing molecular tools for reliable serotyping.

## Results

### Discovery of conserved right-border gene *glpX* of CPS gene cluster

We selected 443 *V. parahaemolyticus* strains from different outbreaks during 2006-2017 from sentinel hospitals of Shenzhen, encompassing 71 serotypes and 40 K-serogroups, for whole genome sequencing. Draft genomes of good quality were obtained, with an average of 193 contigs and average 5.12 Mbp in total length for each genome. The average N50 length and average N90 length of assembled contigs are 335.44 kbp and 59.48 kbp, respectively.

CPS loci gene clusters of total 443 *V. parahaemolyticus* strains were extracted from their whole genome sequences. After assembly, CDS prediction was conducted as described in Methods and S1 Fig. We found that left-border gene *VP0214* (gene symbol: *gmhD*) is conserved in all strains, but the right-border gene *VP0238* (gene symbol: *rig*) and secondary right-border gene *VP0237* are not fully conserved. The distribution frequency of *VP0238* in all 443 strains is 49.7% (52.4% in Fig 1A analysis), in 40 K-serogroups representative strains is 87.5% (see bellowing results). The distribution frequency of *VP0237* in all 443 strains is 86.2% (84.9% in Fig 1A analysis), and in 40 K-serogroups representative strains is 72.5% (See results below). Thus, we inferred that CPS gene cluster of *V. parahaemolyticus* has a more conserved right-border gene. A polysaccharide related gene *glpX* near *VP0238* was found fully conserved in all strains. Therefore, we consider it to be a potential right-border gene and subject it to validation in subsequent analysis. Based on high-quality assembly, *glpX* could be extracted from 418 CPS gene clusters.

**Fig 1.**
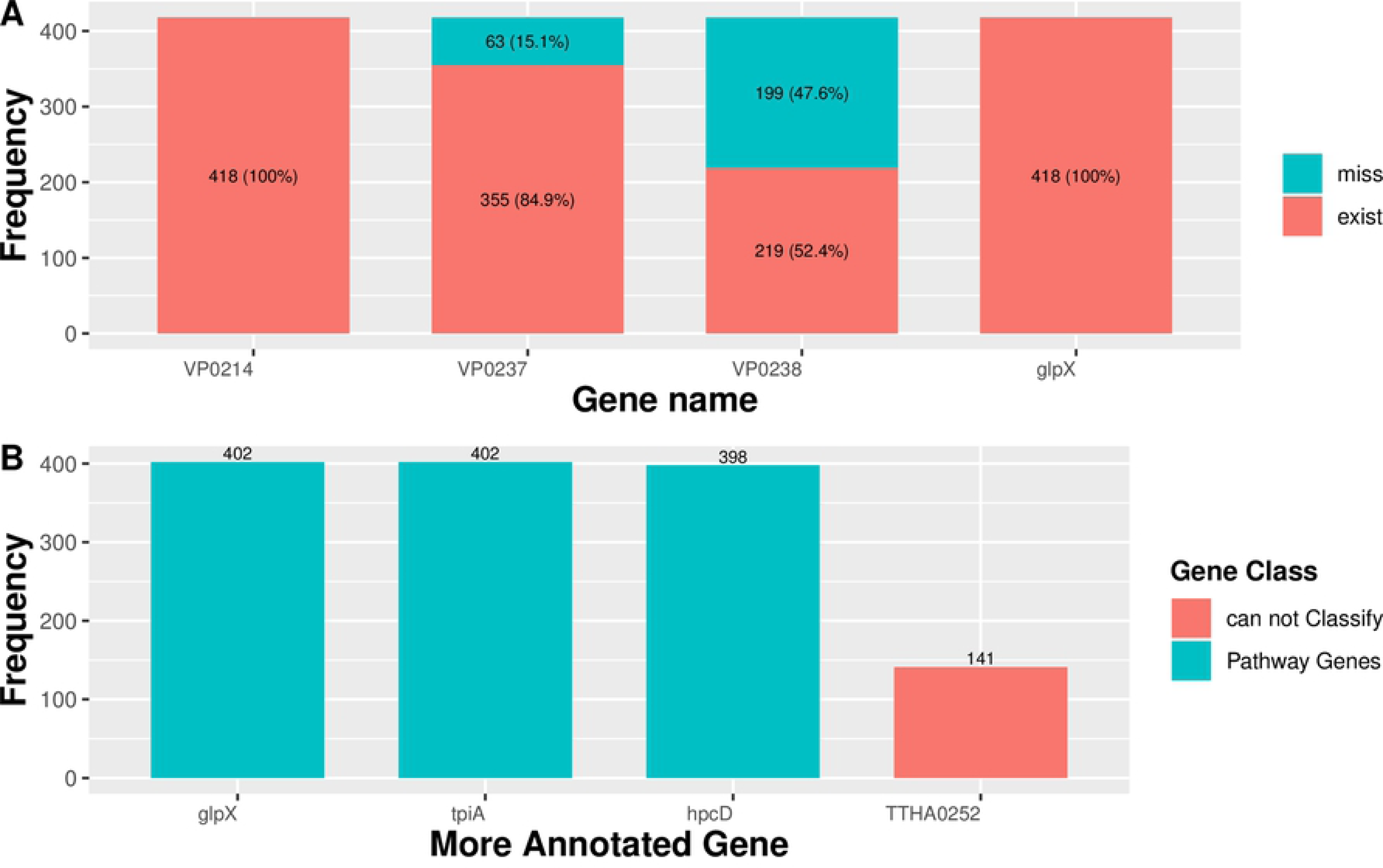
The identification of novel right border gene *glpX* of CPS gene cluster. **(A)** The distribution frequency of previous report border genes and novel right border gene *glpX* in 418 strains *V. parahaemolyticus*, in which intact CPS gene cluster could be assembled. According previous studies [26, 27], *VP0214* is the left border gene of CPS gene cluster of K6 serogroup, *VP0238* is the right border gene, and *VP0237* is the second right border gene. *glpX* is the new right border gene discovered in this study. The blue and pink bars refer to the frequency of gene missing and existing respectively. **(B)** The conserveness of additional genes ORFs between previous right border gene *VP0238* (or *VP0237*) and new right border gene *glpX*. In this analysis, among above mentioned 418 strains, total 402 strains have *VP0238* or *VP0237* in the CPS gene cluster, and generally, 4 additional genes were identified between previous right border gene *VP0238* (or *VP0237*) and new right border gene *glpX*. The bars in figure refer to the distribution of annotated gene group in total 402 strains, additionally, the color of bars represents the gene class of gene group.

In order to determine whether CPS gene cluster of *V. parahaemolyticus* has a conserved right-border gene and ensure inclusion of an entire CPS gene cluster, we check downstream ORFs along *VP0238* one by one and annotated them until no polysaccharide related genes can be found. We found a polysaccharide related gene *glpX* which is about five ORFs downstream of *VP0238*, distributes in all 443 strains including in the 418 well assembly CPS gene clusters (Fig 1A). Furthermore, the right neighboring ORF of *glpX* is *zapB* (UniProt ID: P0AF3, protein: Cell division protein ZapB), which belongs to cell division related gene cluster (data not shown). The genes between *glpX* and *VP0238* are polysaccharide-related genes (Fig 1B): *glpX*, *tpiA* and *hpcD*; more specifically, *tpiA* (402/402 in Fig 1B analysis) and *hpcD* (398/402 in Fig 1B analysis) also are fully conserved in all 418 well-assembled CPS gene clusters (including in 40 K-serogroups representative strains, see Table 1). In summary, *glpX* is an accurate right-border gene of CPS gene cluster in *V. parahaemolyticus*. The following analyses are all based on the entire CPS gene clusters which are extracted by left-border gene *VP0214* and new right-border gene *glpX*.

**Table 1.**
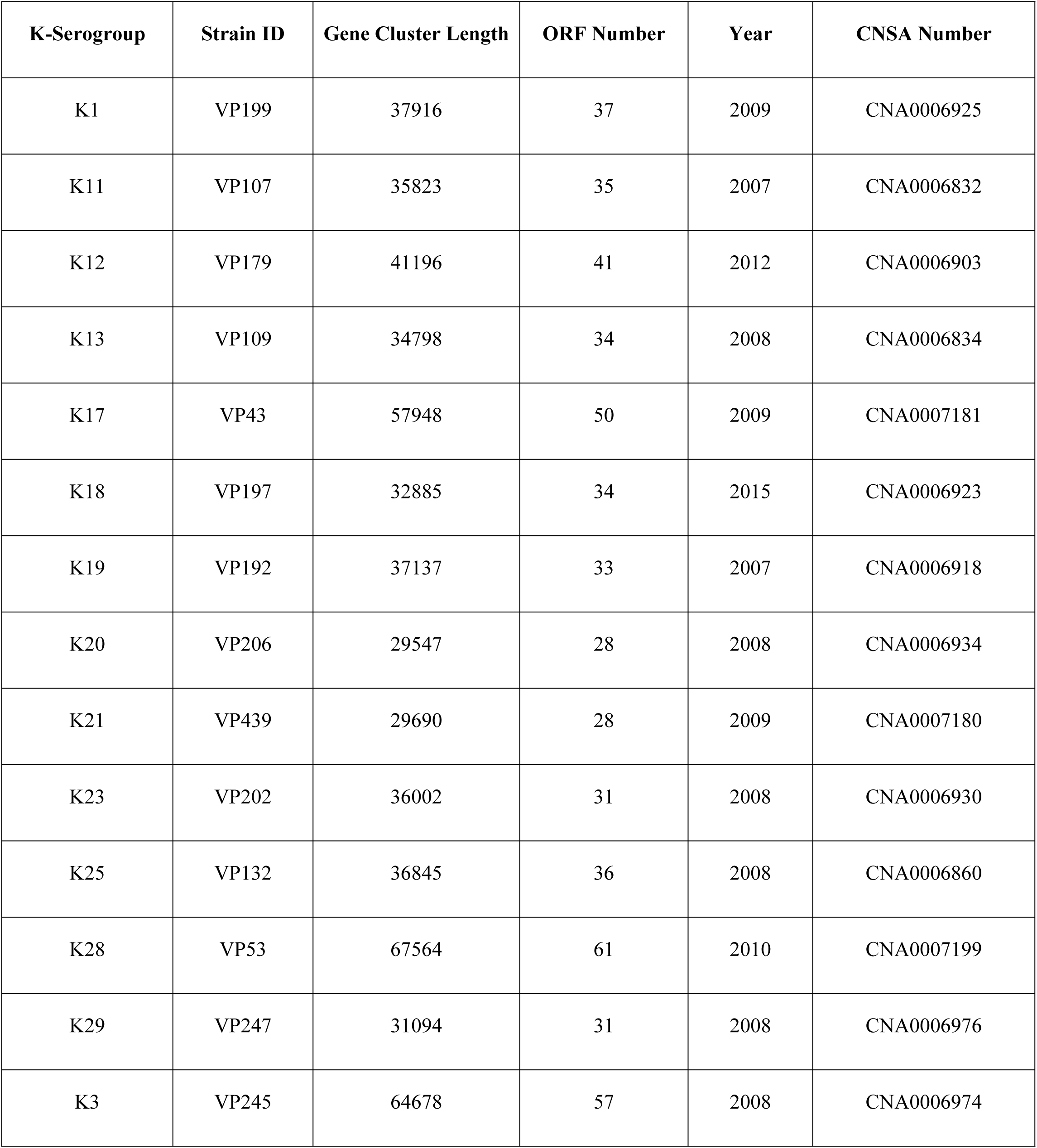
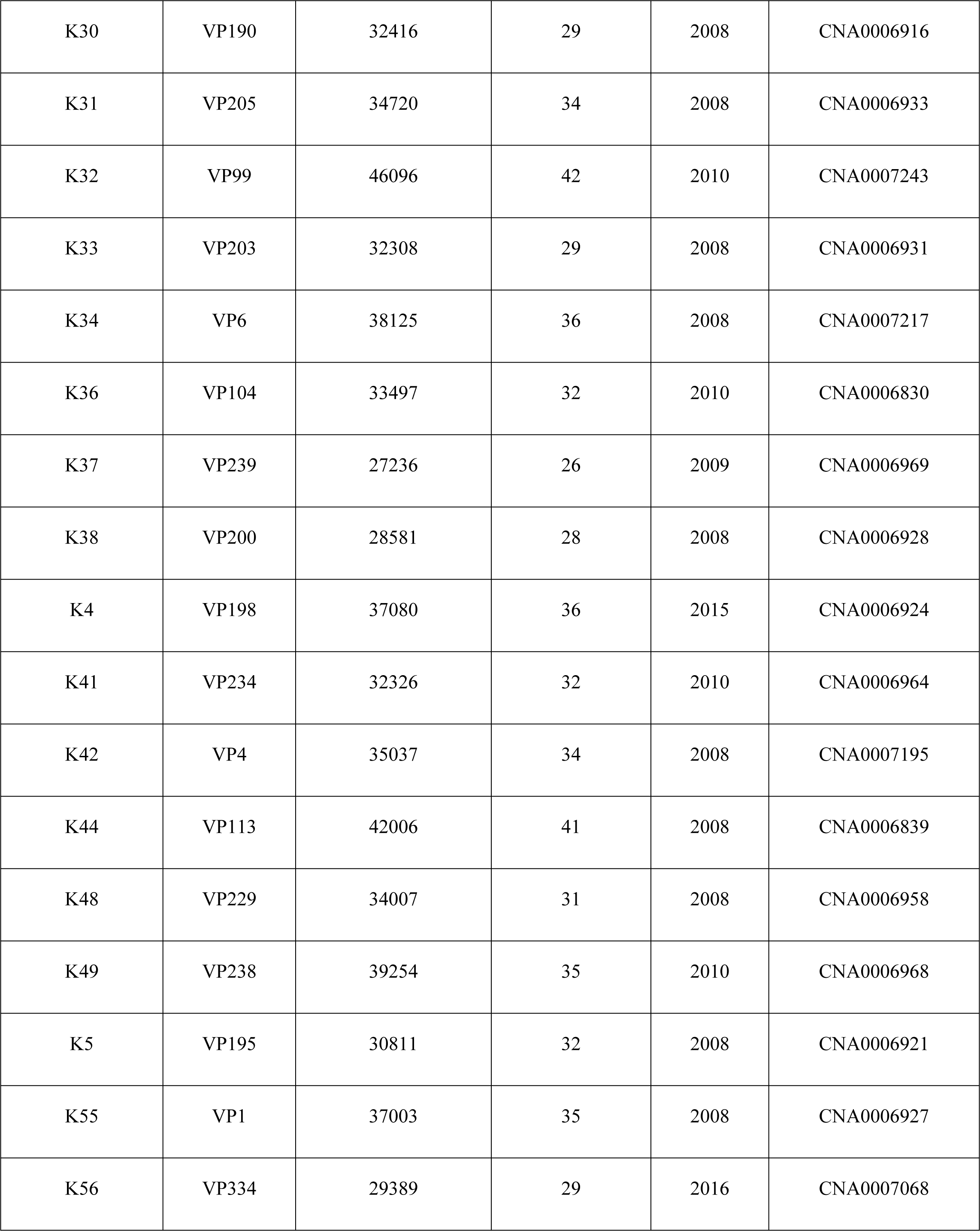
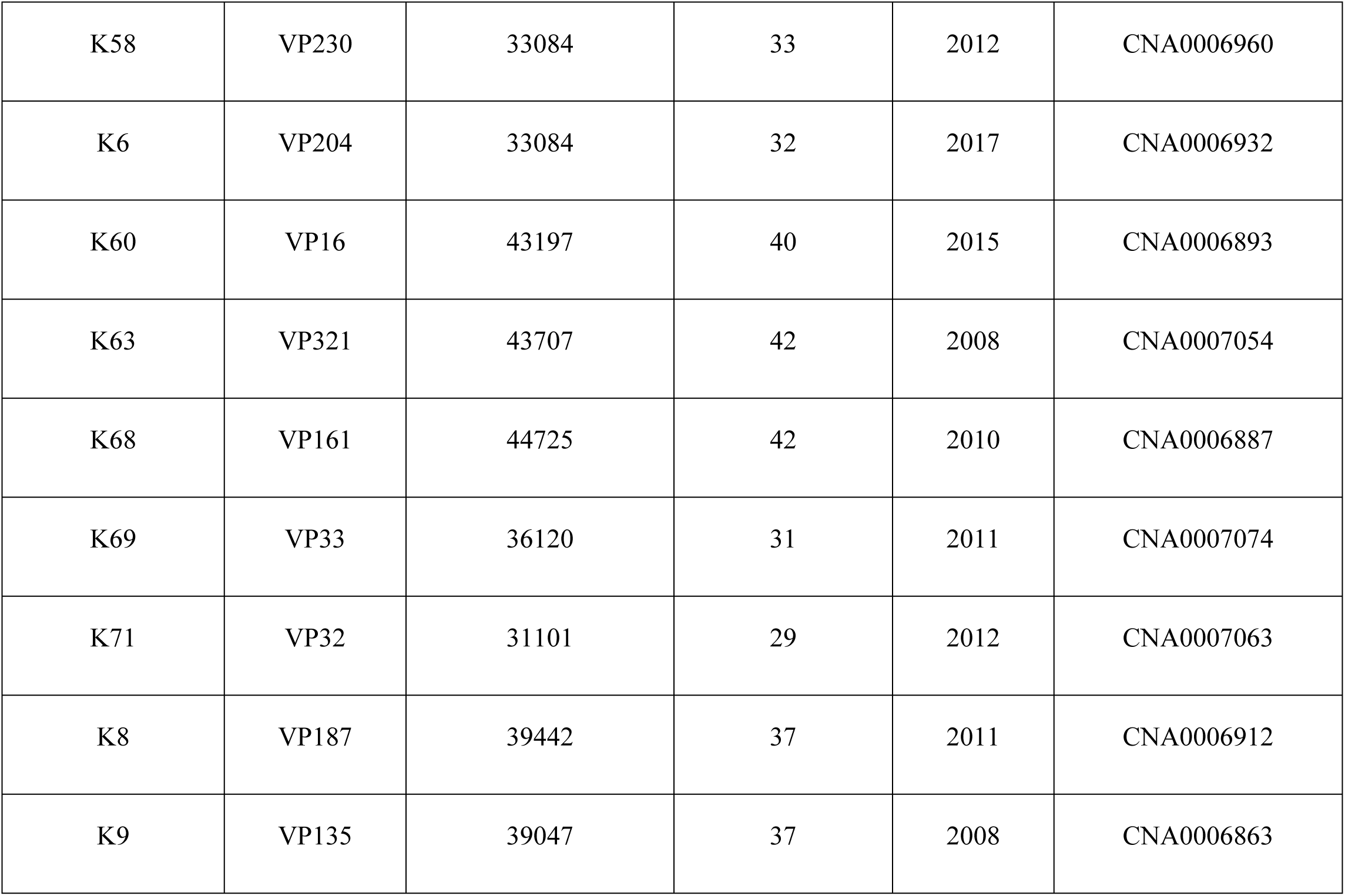
Representative strains of 40 K-serogroups.

### Relationship between serotype/K serogroup and CPS loci length/ORF number

In order to evaluate the variance in coding ability of CPS gene clusters of different serotype/K serogroups, the length/ORF number of CPS loci were calculated and compared. The average length of 40 K-serogroup gene cluster was found to be 37.84 kbp. More specifically, K28 has the longest gene cluster (67.56 kbp), while K38 has the shortest (28.58 kbp). For ORF number, the average of 40 K-serogroup gene cluster is 36. Correspondingly, ORFs are most abundant in K28 (61 ORFs), and least abundant in K38 (28 ORFs) (Fig 2, S2 Fig, S1 Table). Based on this, we deduce that CPS gene cluster length is directly proportional to their ORF number in different K-serogroups.

**Fig 2.**
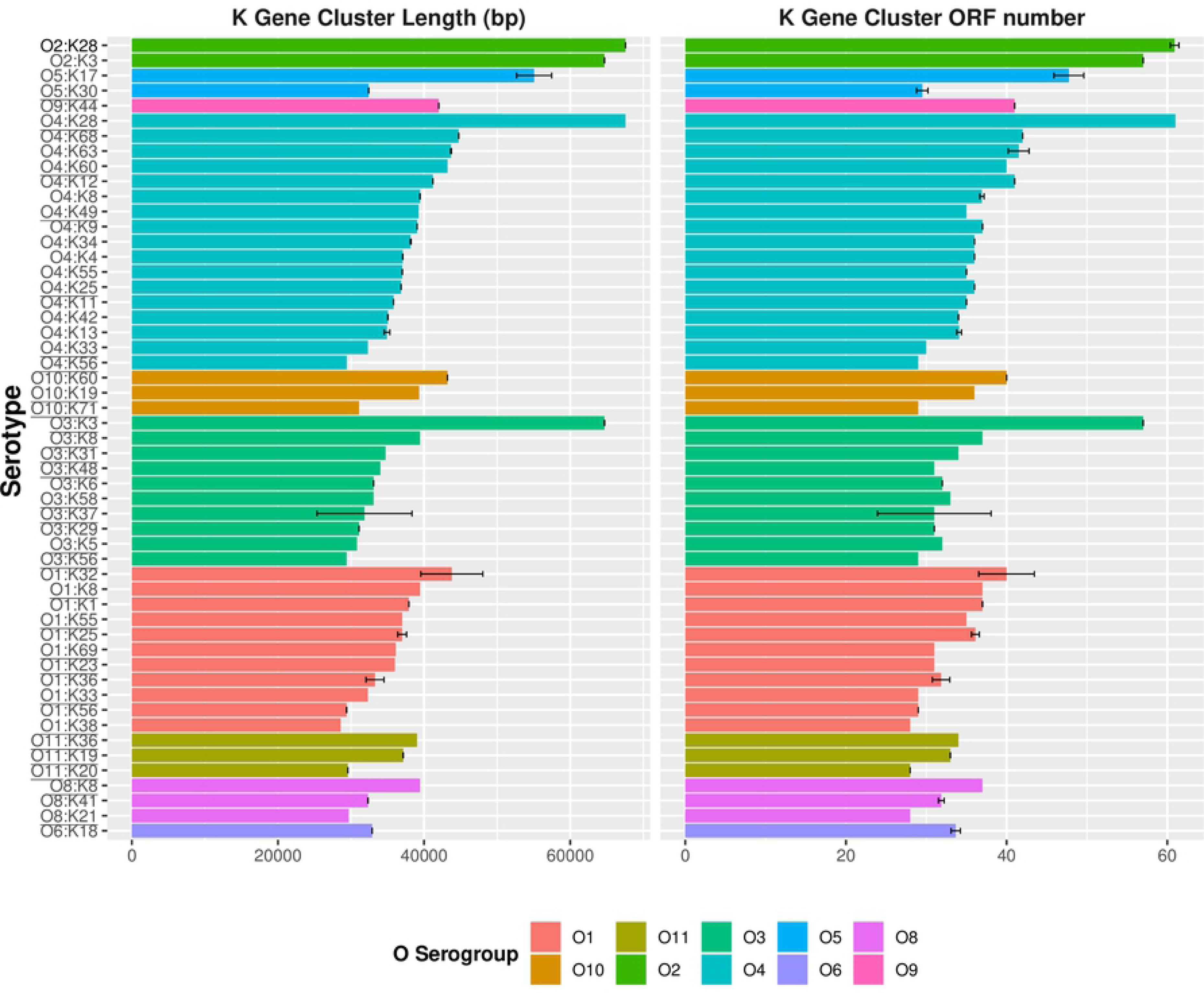
The coding capacity and difference of CPS gene cluster of different serotypes. Bar graph showed the CPS gene cluster length and ORF number distribution of different serotypes without KUT/OUT, and these serotype grouped by O-serogroup with same color. The left bar graph represents the mean of different K-serogroups’ CPS gene cluster length, and the right bar graph represents the mean of ORF number. The error bars give standard deviation for K serogroups which have more than one sample, and underline represents the K-serogroups whose sample number less than 3. The bars are ordered by CPS gene cluster length and ORF number respectively in every O-serogroup.

Interestingly, the lengths of CPS gene cluster from K-serogroups combined with O2 forming serotypes (namely O2:K28 and O2:K3) are the apparently longer than that of K-serogroups combined with other O-serogroups, which is in agreement with the ORF number (Fig 2). This suggests that K28 and K3 have a closer evolutionary relationship. The following evolution analysis indeed supports this view.

### Representation of 40 CPS gene cluster’s structure, function and their differences

In order to clarify coding gene structure and gene function of 40 K-serogroups gene clusters, we selected one representative strain from each K-serogroup to analyze and cluster coding genes within CPS gene cluster (Table 1). Accordingly, 40 K-serogroups’ coding genes were clustered into 219 genes which were subsequently classified into 4 classes, including 48 pathway genes, 12 processing and transportation genes, 6 glycoltransferase genes, and 153 others (4 known functions but cannot be classified, and 149 unknown function) (Table 2).

**Table 2.**
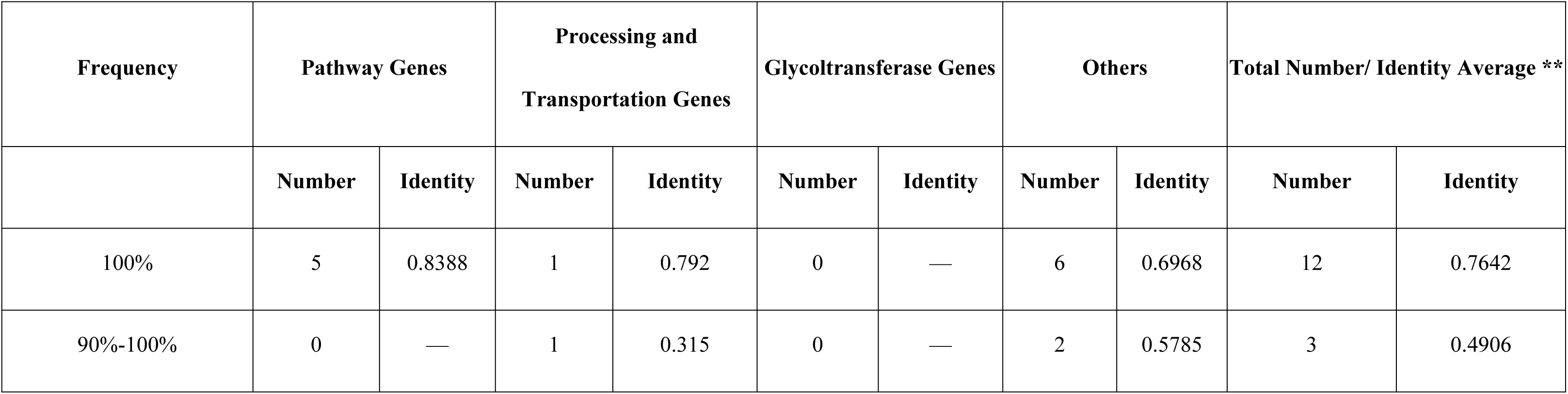
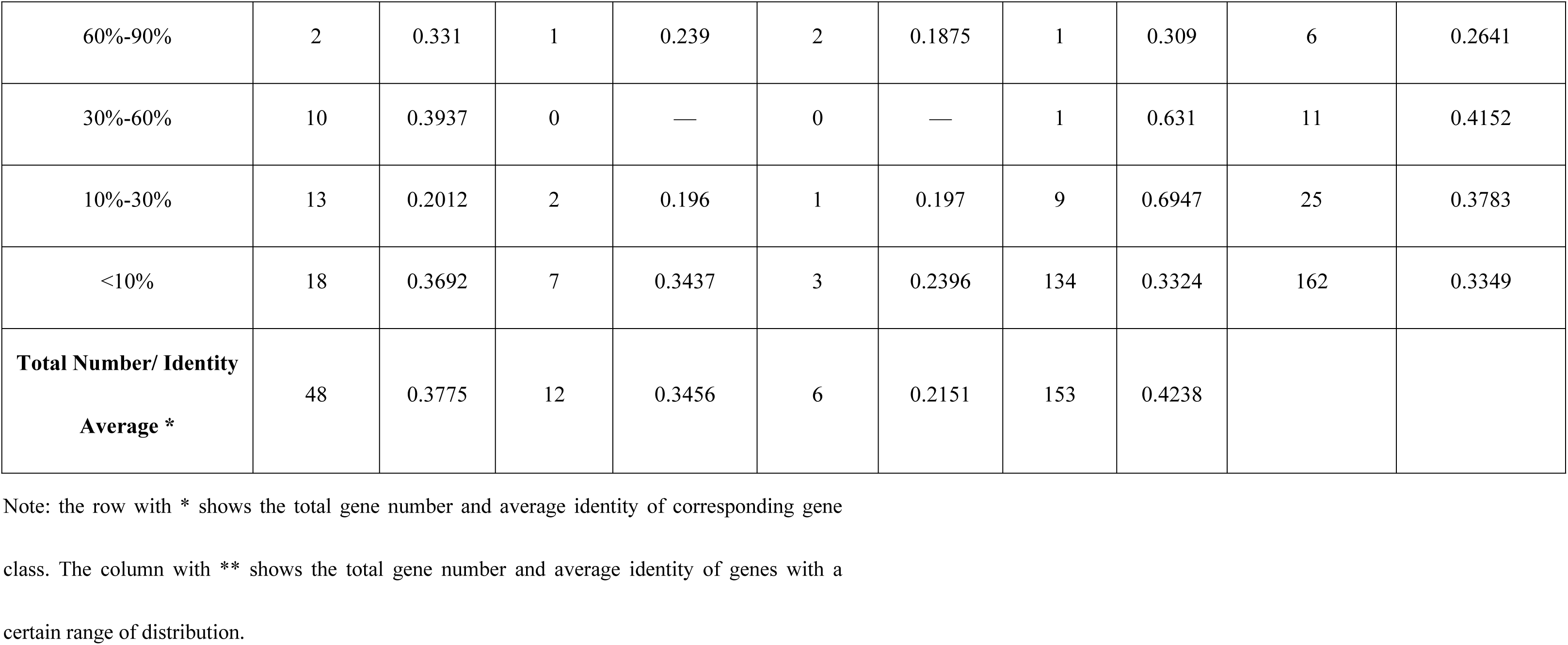
Gene number and identity average of Core-pan gene and gene class.

#### Structural characteristics of CPS gene cluster’s coding region

To characterize the CPS gene cluster further, we plotted the gene structures of 40 K-serogroups according to annotations. In 40 K-serogroups, the genes in the flank region are more conserved than those in the middle region, with a distribution frequencies of more than 90%. More specifically, there are a total of 15 core genes, whose distribution frequency was more than 90%: 12 of them have distribution frequency of up to 100% (within-genes mean pairwise identity is between 55.4%-97.4%). Another 3 genes have a distribution frequency of between 90%-100% (*VP0218*, *VP0219*, *wzc*; within-genes mean pairwise identities are 48.8%, 66.9%, and 31.5%) (<u>S2 Table</u>, Fig 3C). In addition, these core genes are single-copy, and have a structurally conserved order in the flank region of 40 different K-serogroups, with approximately eight core genes except for K3 and K28 (four core genes) located in the left flank region and seven core genes located in the right flank region (Fig 4, <u>S3 Fig</u>).

**Fig 3.**
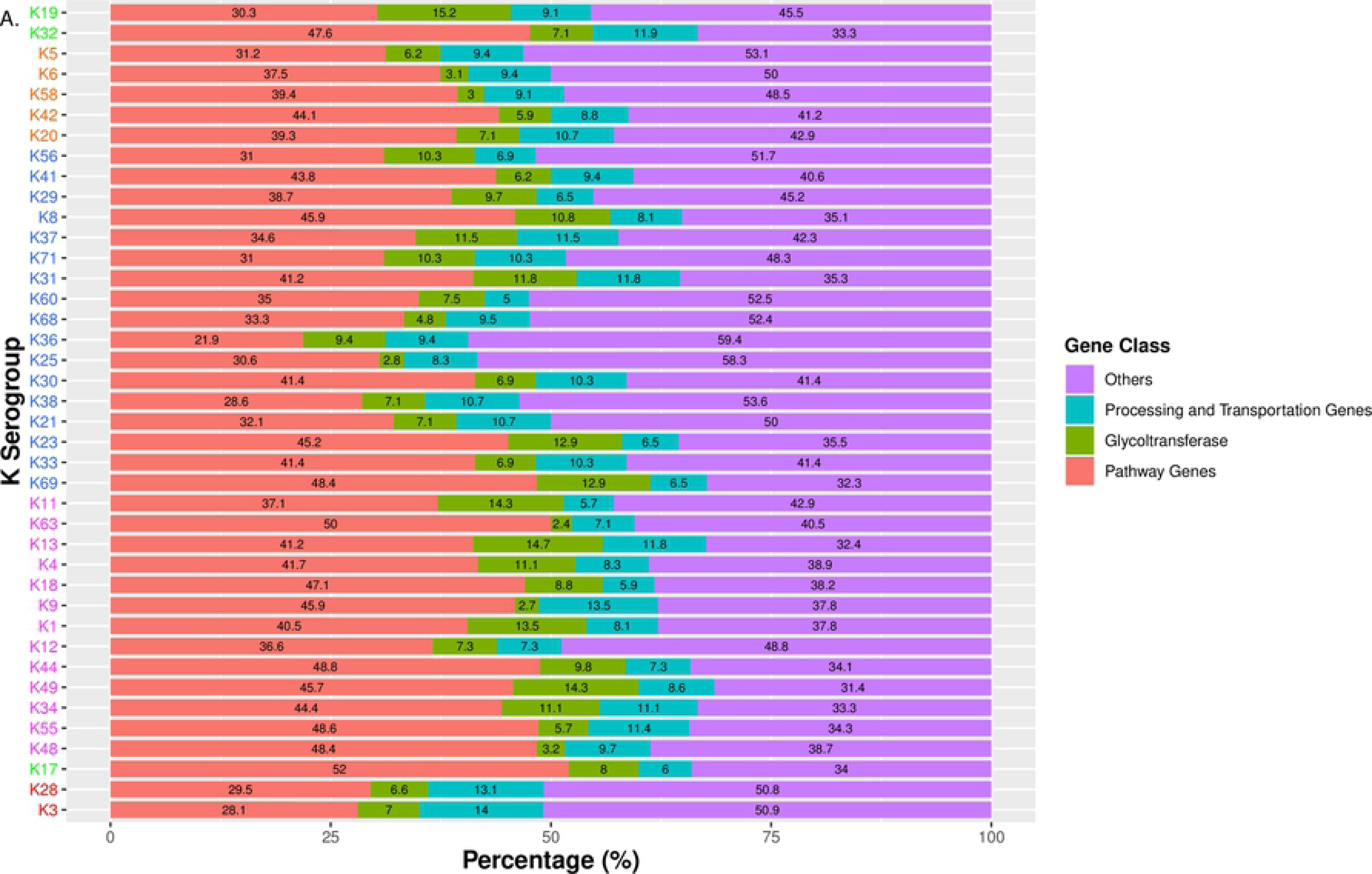
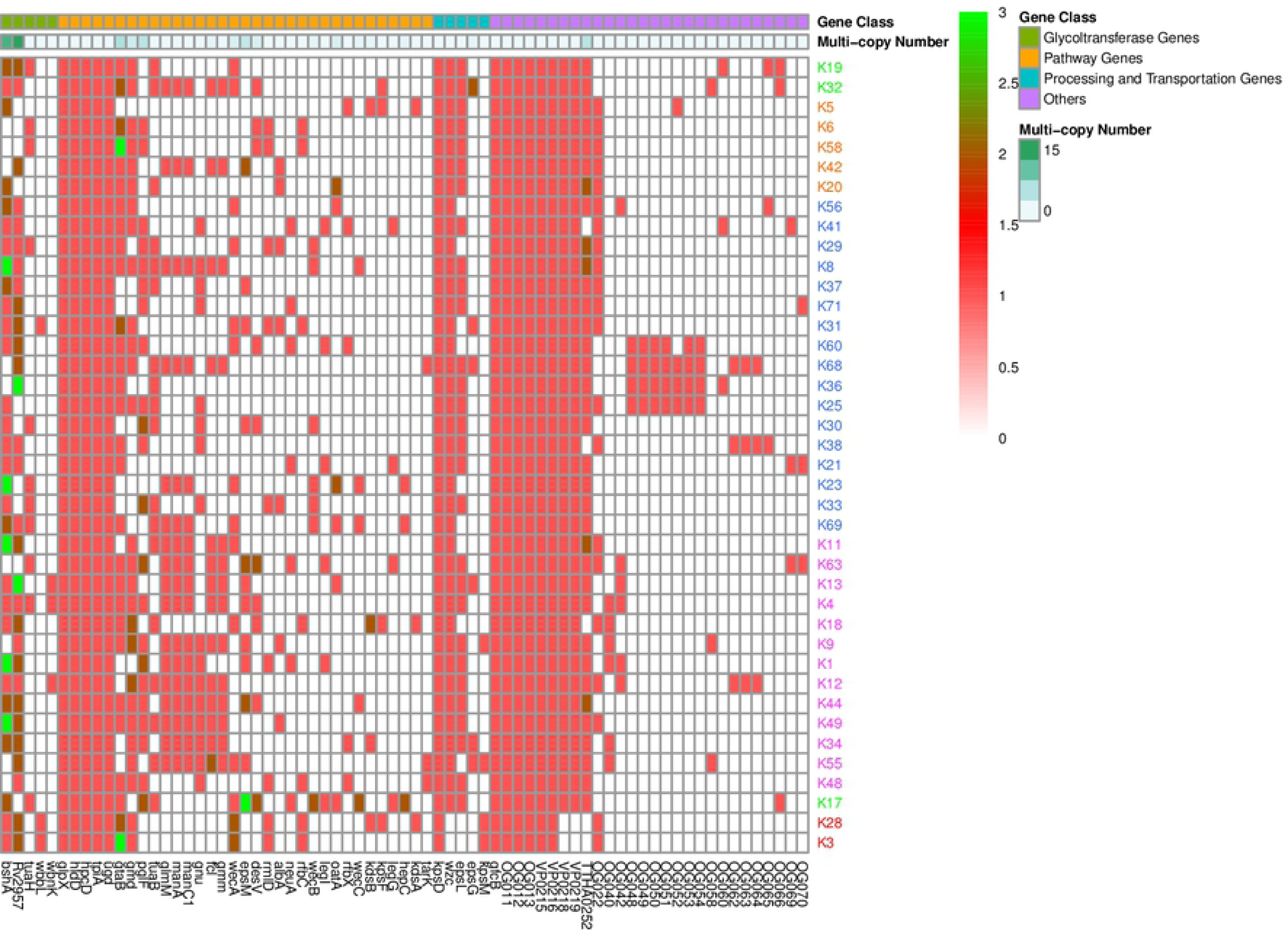
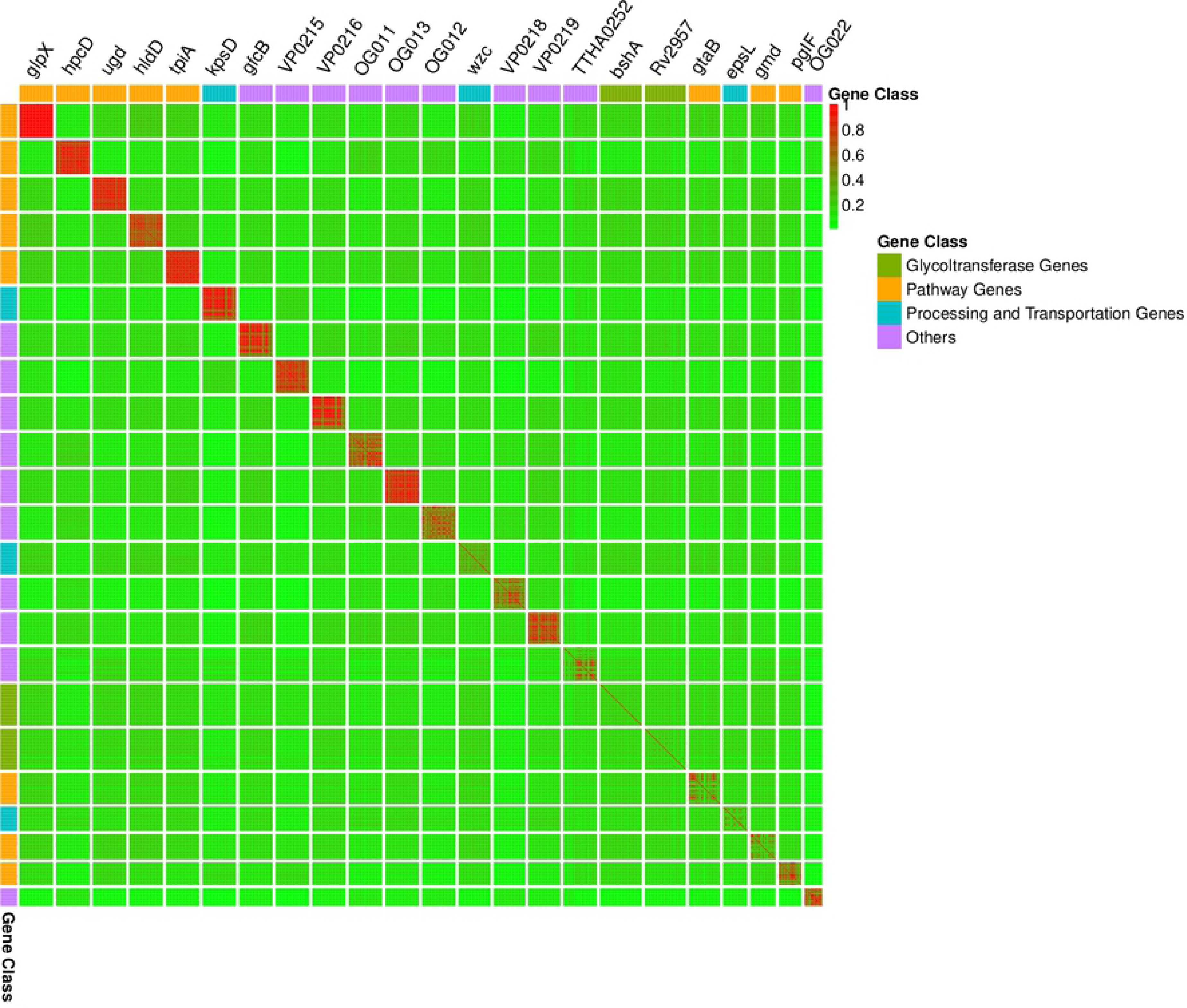
Contents, distribution, diversity of genes coding by 40k. **(A)** Functional contents of 40 K serogroups displayed by proportion of four gene classes. **(B)** Gene presence or absence heatmap of 40 K-serogroups. Columns, corresponding to different genes, were ordered by gene class and occurring frequency in 40 K-serogroups. Those genes existing in less than 2 K-serogroups were not included, for all genes version, please refer S4 Fig. **(C)** Heatmap profile for gene diversity assessment by within-gene pairwise identity. Genes occurring in more than 20 K-serogroups are displayed and sorted by distribution frequency and gene function class. Each cell shows the pairwise identity of all ORFs from two gene groups. In (A) and (B), K-serogroups were classified, ordered and colored classified according Fig. 5 (red: **Group 1**, green: **Group 2**, orange: **Group 3**, blue: **Group 4**, purple: **Group 5**).

**Fig 4.**
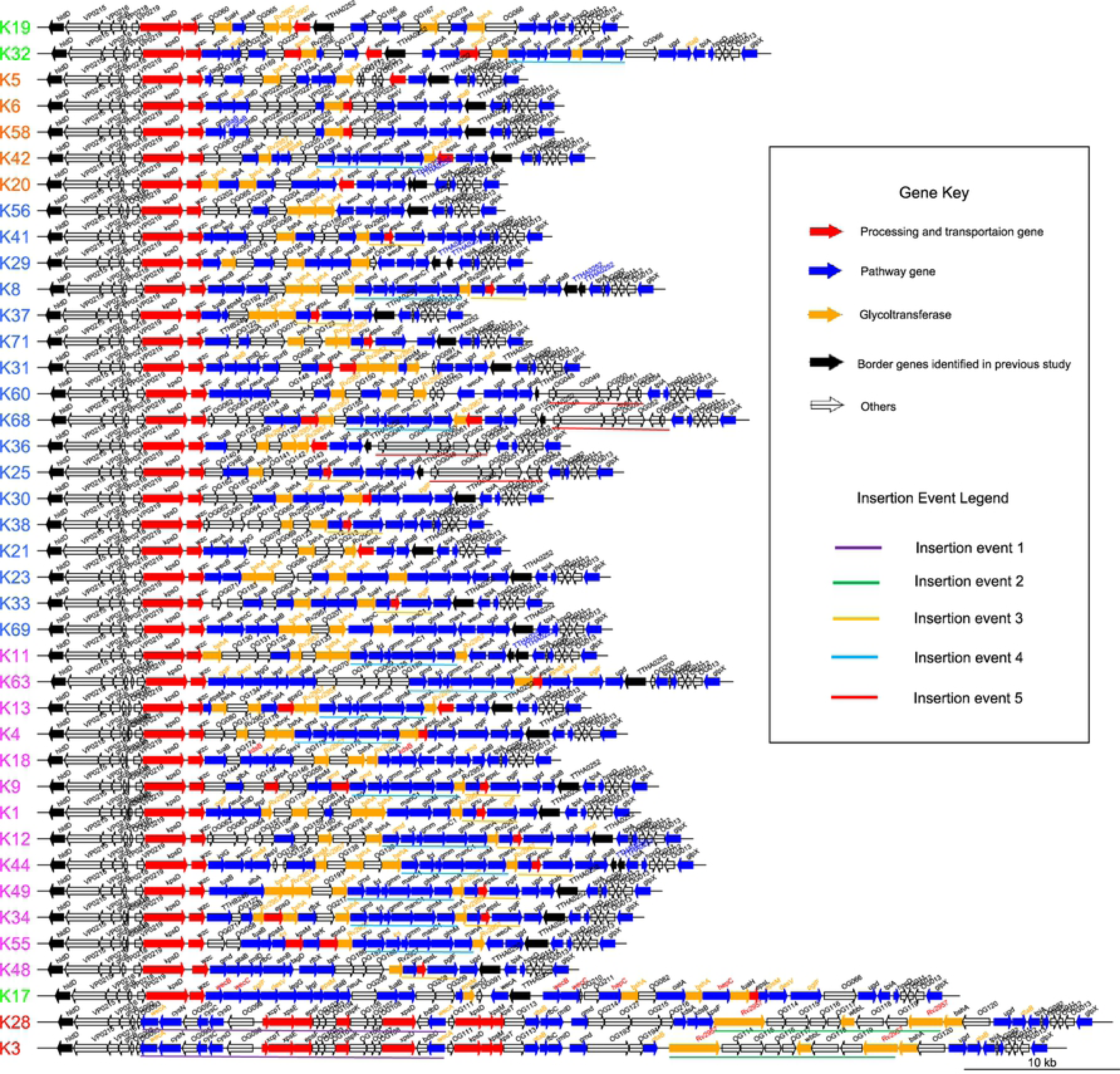
Genetic structure of CPS loci of 40 reprehensive K-serogroups. Every line represents a gene cluster, and one arrow refers to one gene, the direction of arrow refers to the direction of genes. There are four fill colors of arrows represent four gene function classes: red refers to **processing and transportation genes**, blue refers to **pathway genes**, orange refers to **glycotransferase genes**, transparent refers to **others** (gene function unknown or can’t be classified), black refers to two border genes *VP0214* and *VP0238* identified in previous study [27]. In addition, gene names above arrows are assigned 4 colors: black represents genes that didn’t have multi-copies in the corresponding K-serogroup, red represents multi-copies originated by gene self-duplication, blue represents multi-copies originated by nonsense mutation, and orange represents multi-copies originated by recombination. Finally, underlines of five different color highlight five insertion recombination events, and their map relationship is showed in legend. K-serogroups were classified, ordered and colored classified according Fig. 5 (red: **Group 1**, green: **Group 2**, orange: **Group 3**, blue: **Group 4**, purple: **Group 5**).

For the remaining 207 genes, 78% of them are distributed in fewer than three K serotypes (<u>S2 Table, S4 Fig</u>) and are mostly located in the middle regions of the CPS gene cluster (Fig 4). Average within-genes identity is 34.8%, which is less than those in core-genes. Moreover, 16 genes have multi-copies in several K serotypes (Fig 3B, <u>S4 Fig</u>, Table 3). The diversity of gene types and position in the middle region are associated with diversity of the CPS loci in *V. parahaemolyticus*. This is consistent with the CPS gene cluster characteristic among *Escherichia coli* K1, K4, K5, *Neisseria meningitidis* serogroup B, and *Pasteurella multocida* types A, D, and F [31].

**Table 3.**
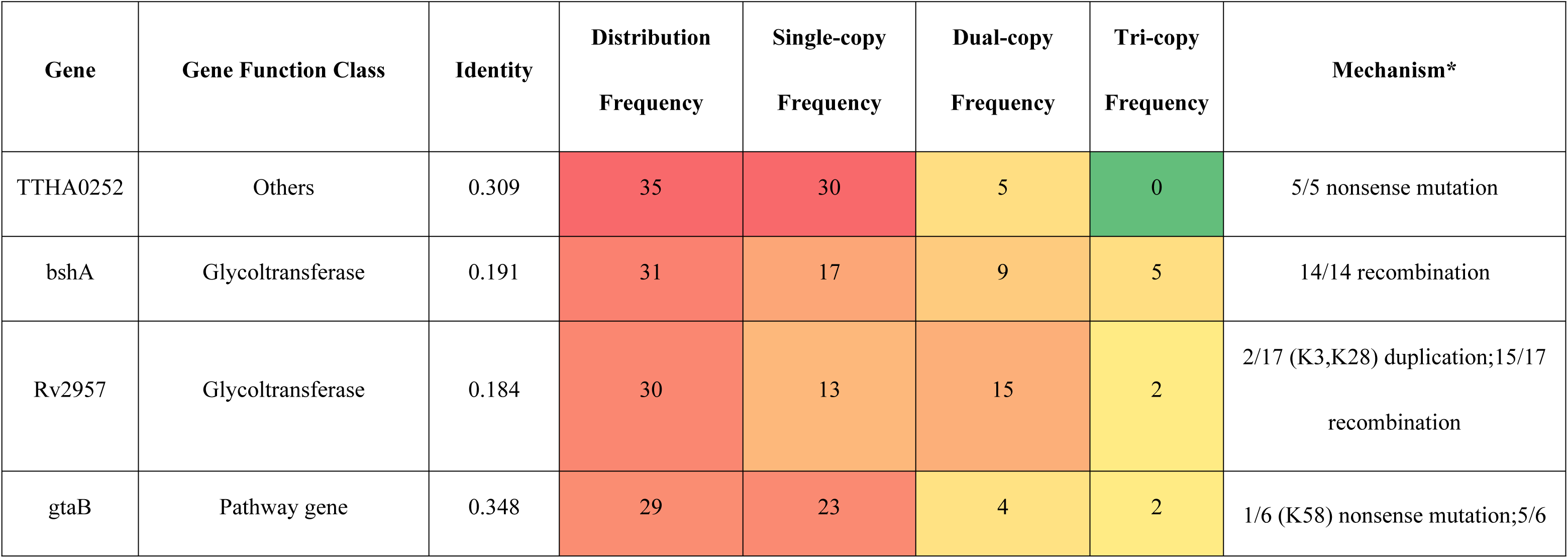
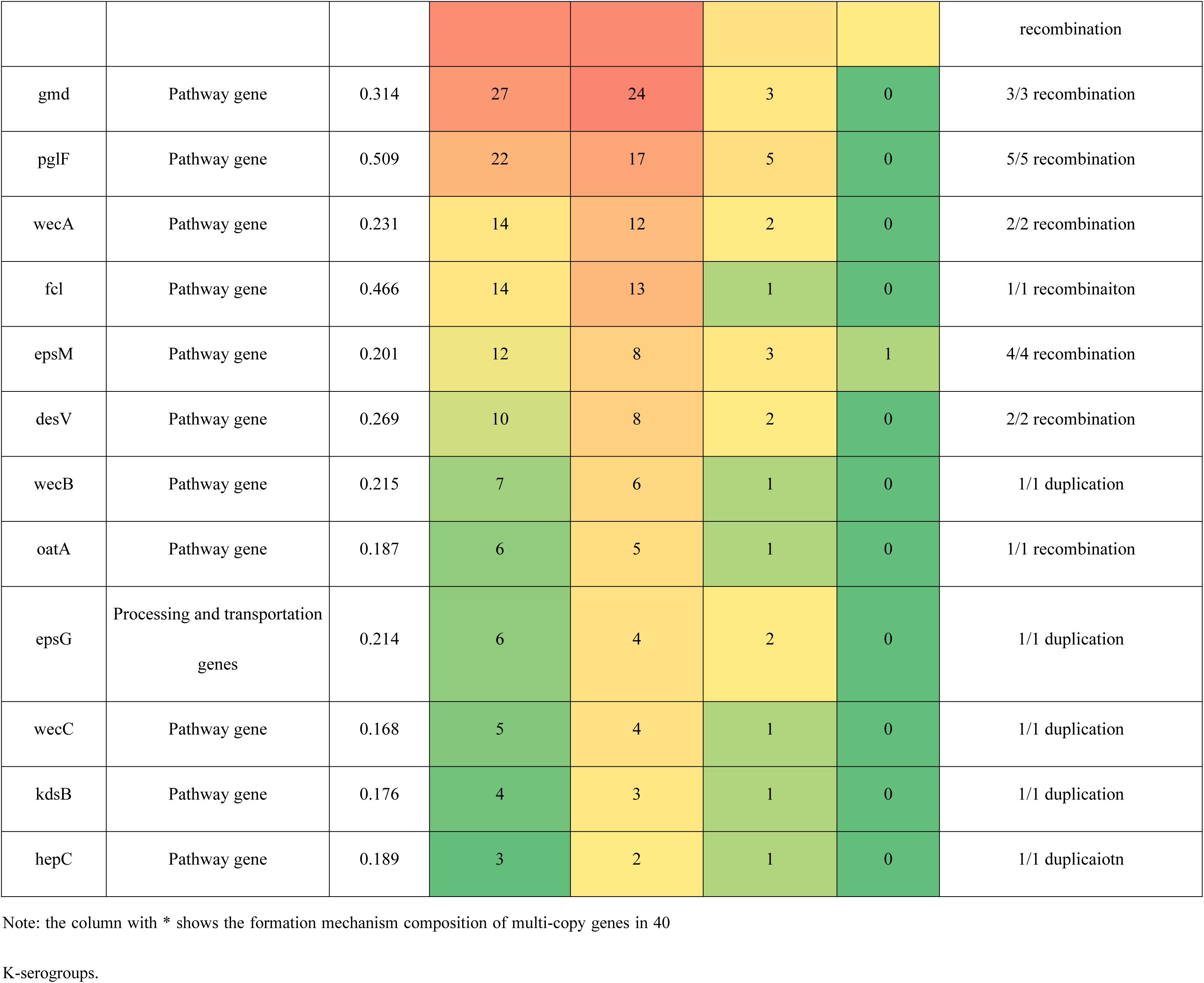
Composition and frequency of multi-copy genes.

In addition, genes whose coding direction is from right to left are mostly located in the flank region of CPS gene cluster, while genes whose coding direction is from left to right are mostly located in the middle region. In the flank region, there are 10 genes (left flank region: *hldD*, *VP0215*, *VP0216*, *gfcB*, *VP0218*. the right flank region, *glpX*, *OG013*, *OG012*, *tpiA*, *TTAA0252*) coding from right to left are conserved in 40 K-serogroups. However, the genes in middle region mostly code from left to right except for K28 and K3. Notably, in K28 and K3, there are more continuous genes coding from right to left, which is distinct from other K-serogroups (Fig 4). The following insertion sequence analysis indicates that these ORFs coding from right to left arose from 2 insertion events.

#### Functional characteristics of CPS gene cluster’s coding region

##### Pathway genes

Pathway gene class, containing 48 genes, are the largest in 3 gene function classes (Table 2), and are mainly located in the middle and right flank region of CPS gene clusters (Fig 4). The average percentage of pathway gene class in each K-serogroup is up to 39.48%: more specifically, K17 has the largest (52%) while K36 has the least pathway genes percentage (21.9%) (Fig 3A). There are total 5 genes (*glpX*, *hldD*, *hpcD*, *tpiA*, *ugd*) whose distribution frequency in 40 K-serogroups is up to 100% in pathway gene class, which is more than other 2 gene classes, what’s more, sequences of these core genes in 40 K-serogroups are highly conserved than any other genes (the average of their within identity is 83.9%) (Table 2, Fig 3B, Fig 3C). Some of pathway genes are reported in other enteropathogenic bacteria responsible for biosynthesis precursor of CPS. *gtaB* and *glmM*, encoding UTP-glucose-1-puridylyltransferase and phosphoglucosamine mutase respectively, exist in 72.5% and 42.5% of 40 K-serogroups (<u>S2 Table</u>), and were reported existing in many pathogenic bacteria, play roles in producing two CPS precursor, namely UDP-glucose and UDP-*N*-acetylglucosamine [31]. In addition, *ugd* encoding UDP-glucose dehydrogenase, exits in all 40 K-serogroups, which also reported in play roles in the biosynthetic pathway for CPS in *Escherichia coli* K4 and K5[31]. Two genes with minor frequency, *wecB* (20%) and *neuA* (17.5%) encoding UDP-*N*-acetylglucosamine-2-epimerase and CMP-N-acetylneuraminate cytidylyltransferase were reported for biosynthesis polysialic acid in *Escherichia coli* K1 (S2 Table). In summary, the core pathway genes of 40 K-serogroups might be essential for synthesis of the common precursor of CPS, while the pan pathway genes maybe catalyze the common precursor to different product, which will change the chemical properties and immunogenicity of CPS.

##### Processing and transportation genes

Processing and transportation genes are responsible for forming the CPS repeat units and translocating mature CPS to cell surface subsequently. [32]. Totally 12 genes were identified to be processing and transportation genes class, and mainly located at the left flank region of CPS gene clusters. The average percentage of this gene function class in 40 K-serogroups is 9.23%: K13 and K31 have the most (11.8%) while K60 has the least processing and transportation genes percentage (5%) (Fig 3A). Among them, two core genes (*kpsD* and *wzc*) were identified: *kpsD* gene, encoding polysialic acid transport protein (UniProt ID: Q03961), exists in all 40 K-serogroups with higher sequence identity 79.2%; while *wzc* gene, encoding Tyrosine-protein kinase (UniProt ID: P76387), exits in all 40 K-serogroups with higher sequence identity, except K28 and K3 with low sequence identity 31.5% (<u>S2 Table</u>, Fig 3B, Fig 3C). According to the annotation result of UniProt database, *kpsD* is involved in the translocation of the polysialic acid capsule acorss the outer membrane to the cell surface, and *wzc* probably invovled in the export of colanic acid from the cytosome to outer membrane, thus we speculated that *kpsD* and *wzc* can non-specifically process and transport very distinct CPS structure. According to a recent study by Olaya Rendueles *et al.* [33], CPS can be biosynthesized by using one of the following five mechanisms recognized by processing and transportation genes: Wzx/Wzy-dependent, ABC-dependent, synthase-dependent, PGA and Group IV. In our study, K28 and K3 belongs to an ABC-dependent mechanism by our annotation strategy, while other 38 K-serogroups cannot be identified by this approach.

##### Glycoltransferase genes

6 genes can be classified into glycoltransferase genes class, and are mainly located in the middle region of CPS gene clusters (in orange in Fig 4). The average proportion of glycoltransferase genes in 40 K-serogroups is the least which is equal to 8.43% in three gene classes: K19 has the most (15.2%) while K63 has the least percentage 2.4% (Fig 3A). These genes all distributed with less frequencies among 40 K-serogroups, ranging from 5% to 77.5%, and at sequence levels with lower within pairwise gene identity (the average of them within identity is 21.5%) (Table 2, Fig 3B, Fig 3C). The diversity and uniqueness among K-serogroups of glycoltransferase genes might lead to the position and kind of glycosyl that polysaccharide combined different, which cause the diversity of CPS [34].

### Evolution analysis of 40 K-serogroups CPS gene cluster

In order to investigate the evolutionary relationship of 40 K-serogroups, a species tree was constructed by using Orthofinder and was rooted at K28 and K3 according to the phylogenetic relationship of core genes and the genetic structure of CPS locus (Fig 5, <u>S5 Fig</u>). We found frequent recombination took place by sequence insertion during the *V. parahaemolyticus* CPS locus evolution, consistent with previous studies in other pathogenic bacteria [33, 35–37]. Based on the phylogenetic relationship and genetic structure, a 40 K-serogroups evolution tree can be divided into totally 5 groups (group1-5) with each being characterized by certain insertion events (Fig 5).

**Fig 5.**
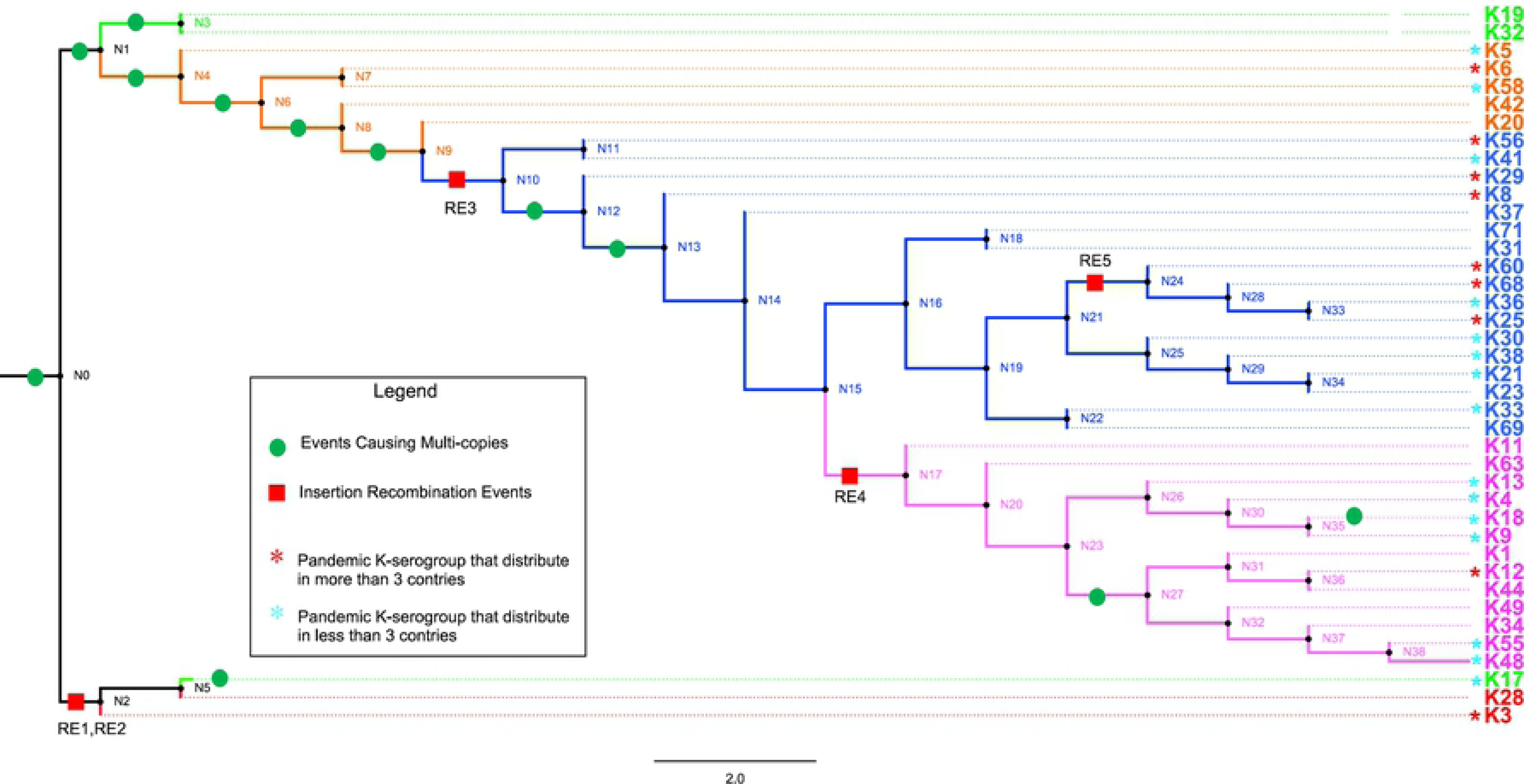
The evolution of 40 K-serogroups. This figure shows the divergence process and evolution relationship of 40 K-serogroups. Five groups of 40 K-serogroups were indicated by different colors for the branch and tips: red refers to **Group 1**, green refers to **Group 2**, orange refers to **Group 3**, blue refers to **Group 4**, and purple refer to **Group 5**. Node ID were displayed, and the red square and green circle on certain branches refer to that insertion events and the duplication events. For gene duplication events, the concrete information is listed in Table 4. For insertion events, the sequence of **RE1** (insertion event 1) contains *wecA*, *cysN*, OG095, *cysC*, *cysD*, OG098, *xcpT*, *xpsE*, *epsF*, *OG102*, *OG103*, *OG104*, *gspK*, *OG106*, *OG107*, *OG108*, *xpsD*, *bdbD* and *wecA* (19 genes in total), **RE2** contains *Rv2957*, *OG114*, *OG115*, *OG116*, *OG117*, *wbbL*, *OG118*, *OG119* and *Rv2957* (9 genes), **RE3** contains *gnu*, *espL* and *pglF* (3 genes), **RE4** contains *gmd*, *fcl*, *gmm*, *mand*, *glmM* and *manA* (6 genes), **RE5** contains *OG048*, *OG049*, *OG050*, *OG051*, *OG052*, *OG053* and *OG054* (7 genes). Pandemic K-serogroups were indicated by star (*) according previous study [21].

**Table 4.** Distribution of duplicaiton events on species tree.

|  | Node | Event | Total Event |
| --- | --- | --- | --- |
| 1 | N0 | bshA*5, Rv2957*5, gtaB*3, gmd*1 | 14 |
| 2 | N1 | bshA*1, epsM*3, desV*1, wecB*1 | 6 |
| 3 | N3 | Rv2957*1, gtaB*1, wecA*1 | 3 |
| 4 | N4 | bshA*1, TTHA0252*2 | 3 |
| 5 | N6 | gtaB*1 | 1 |
| 6 | N8 | fcl*1 | 1 |
| 7 | N9 | oatA*1 | 1 |
| 8 | N12 | pglF*1 | 1 |
| 9 | N13 | Rv2957*1 | 1 |
| 10 | N27 | Rv2957*1 | 1 |
| 11 | VP43 | wecC*1, hepC*1 | 2 |
| 12 | VP197 | kdsB*1 | 1 |
| **Total** |  | 35 | 35 |

#### Group1

As shown in the above analysis, CPS loci of K28, K3 are much longer and possess more ORFs than do other K-serogroups (Fig 2). They share similar structure and gene class composition proportion except for the 34th-41st and 34th-37th genes from left flank (Fig 4, S3 Fig), but very different from other K-serogroups. As we can see in S5 Fig, K28 and K3 cluster in a clade distinct from others both in core-gene tree and species tree. Thus, K28 and K3 are designated as Group1.

Additionally, we proposed that during the evolutionary process, K28 and K3 were generated by two insertion events from their ancestor, which have similar genetic structure with other K serogroups (Fig 4, Fig 5). Both donor sequences of insertion event 1 and 2 are terminated by the same gene, *wecA* and Rv2957, respectively. The genes in these two insertion sequences uniquely are distributed within group 1, supporting the notion that they were recombined from other species.

To determine the potential donor sources of insertion sequence, we extracted these sequences and blasted them in Genbank. Most hits are from *V. parahaemolyticus*, which are deduced to be homological with our search sequences but not a real donor source. Thus, we focused on the no-*V. parahaemolyticus* hits to identify the donor species. For sequence of insertion event 1, apart from five near full-length matching hits from potential K3 and K28 *V. parahaemolyticus*, six hits come from the same region, located in cysD gene, of other *Vibrio* species, namely *Vibrio crassostreae* (identity 81.6%, coverage 4%), *Vibrio breoganii* (identity 81.6%, coverage 4%) and *Vibrio vulnificus* (identity 80.9%, coverage 4%) (<u>S6 Fig</u>). For event 2, all 3 hits are from *V. parahaemolyticus* (<u>S4 Table</u>). We reasoned that the most recent donor sources of the whole insertion sequences in event 1 and 2 is unidentifiable by the current approach because of low availability of public sequences relevant to the analysis, though we can find that the homologous cysD from the other *Vibrio* species.

#### Group 2

Group2 including K19, K32, and K17 are adjacent in the species tree and form a distinct clade in the core-gene tree at the root (<u>S5 Fig</u>). An important difference between group 2 and others is that: the old border gene *TTHA0252* (*VP0238*) locate at the middle of group 2 gene cluster, while locate at the right flank or absent in other groups. This further support the fact that *TTHA0252* (*VP0238*) is not conserved in *V. parahaemolyticus* CPS gene cluster, and if regard it as right-border gene, will lead to error in extracting gene cluster. In addition, there is a group-specific gene *OG066* at the right flank in group 2.

#### Group 3

Group 3 contains five K serotypes, namely, K5, K6, K58, K20 and K42. This group differentiated from group 2 at node 1, and developed into group 4 and group 5 later (Fig 5), they share similar cluster lengths ranging from 30 kb to 40 kb.

In this group, it is noteworthy that the genetic structure and gene content of K6 and K58 CPS loci are very similar, while the major difference is that the tenth gene *gtaB* (EC 2.7.7.9, protein name: UTP-glucose-1-phosphate uridylyltransferase) from left flank region were separated into two ORFs by the 50^th^ codon mutation into a termination codon. More specifically, the 50^th^ codon of *gtaB* (total 292 codons) in K58 mutated to a termination codon leading to a new ORF of 50 codons, and the 139^th^ codon mutated to an initiation codon leading to another new ORF of 154 codons. O3:K58 is a pandemic serotype which was identified in the post-pandemic period [38]. The high similarity of CPS gene cluster between K6 and K58 indicates that K58 may have originated from K6 by *gtaB* nonsense mutations.

#### Group 4

We proposed that an insertion event 3 has occurred in the ancestor (N10) of group4 and group5 (labeled with bold orange lines in Fig 4, Fig 5). The sequence of insertion event 3 contains three genes *gnu*, *epsL* and *pglF* with conserved order. This insertion sequence specifically exists in group 4 (7/17) and group 5 (7/13) but not in others groups. Blastn search against Genbank found that two hits from *Vibrio alginolyticus* have 98%, 100% coverage and 91.7%, 91.3% identity, which are presumably the potential source of insertion event 3 sequence. In addition, this sequence is also found in many other *Vibrio* species with different coverage and identity such as *Vibrio campbellii*, *Vibrio neocaledonicus*, *Vibrio owensii*, *Vibrio harveyi* and so forth (<u>S4 Table, S6 Fig</u>), indicating that the insertion sequence was promiscuously transferred among *Vibrio* species.

In addition, another more recent insertion event has occurred in K60, K68, K36, K25 located between an old border gene *TTHA0252* (*VP0238*) and *tpiA* destroying most region of *TTHA0252* (*VP0238*) gene, named insertion event 5 (labeled with bold red lines in Fig 4, Fig 5). This is consistent with the insertion sequence in the right junction region of putative CPS gene clusters of K68 and K25 found by Masatoshi Okura *et al.* [26], and the right flank 3 genes, *OG052*, *OG053*, *OG054*, were annotated as transposase genes. We found that the insertion sequences of K68, K36 and K25 were acquired from the same donor source. In K60, however, it was acquired from another donor source. The insertion sequence in K68, K36, and K25 contains 7 genes (*OG048*, *OG049*, *OG050*, *OG051*, *OG052*, *OG053*, *OG054*) that share closer evolutionary relationship in gene trees, whereas in K60, it lacks gene *OG052* (Fig 4). The average identity of insertion sequence in K68, K36 and K25 is up to 99.9% (S7 Fig). More specifically, there are only 4 nucleotide variations in the entire 7234 bp sequence (data not shown), suggesting that the insertion putatively comes from the same donor species and occurs at similar time in recent. Blastn search against Genbank shows that best hits are *Vibrio splendidus* and *Vibrio coralliilyticus* with a coverage of up to 59% and 61%, respectively. The homologous sequence in *Vibrio splendidus* has 59% coverage and 89.9% identity which contains 3 genes *OG048*, *OG049*, and *OG050*. However, the sequence in *Vibrio caralliilytics* has 61% coverage and 94.6% identity, which loses *OG049* and *OG050* compared to the insertion sequence in *V. parahaemolyticus*. In addition, e value of 96 hits are equal to 0 and coverage less than 50%. Interstingly, these hits correspond to the right flank 3 genes which are annotated as transposase genes and come from many *Vibrio* species such as *Vibrio cholera*, *Vibrio vulnificu*s, *Vibrio anguillarum et al.* (<u>S4 Table</u>, <u>S6 Fig</u>). Collectively, these suggest that the sequences of insertion event5 are actively transmitted among *Vibrio* species, which is especially the case for the three transposase genes, *OG052*, *OG053* and *OG054*.

#### Group 5

13 K-serogroups forming an independent clade were classified as group 5 (Fig 5). We inferred that the co-ancestor N17 of these K underwent an insertion event 4 (labeled with blue bold lines in Fig 4; Fig 5), which makes the descendants’ K gene clusters share a six-gene sequence specifically within this group (Fig 5). The insertion sequence contains 6 conserved genes, with a fixed order (Fig 4). The entire insertion sequence exists in group 5 with high frequency (10/13 in this group, mean pair-wise identity 70%), but with low frequencies in group 3 (1/5), group 4 (2/17). We speculate that the insertion sequence might have originated from an insertion events between the ancestral receptor *V. parahaemolyticus* K-serogroup and the donor species, and remained in group 5 during evolution, whereas a gradual loss has taken place in some K-serogroups. Blastn reveals that *Vibrio alginolyticus* (99% coverage, 96.3% identity) and *Vibrio harveyi* (89% coverage, 96 % identity) are the potential donor species. Part of this sequence also was found in *Vibrio alfacsensis* and *Vibrio campbellii* (<u>S4 Table</u>, <u>S6 Fig</u>).

#### Gene duplication driving the evolution of *V. parahaemolyticus* CPS loci

Gene duplication is a mechanism underpinning the generation of new genes and protein functional diversity [39–41]. We found that gene duplication is an important mechanism driving the evolution of *V. parahaemolyticus* CPS loci. As discussed above, in 40 K-serogroups, there are 16 multi-copy genes, most of which being pathway genes (12), whereas others include 2 glycoltransferase genes, 1 processing and transportation gene, and 1 unclassified gene. Remarkably, among these genes, the mean pairwise identities are negatively correlated with the frequency of multi-copy (r=0.5748), especially for genes distributed in more than 10 K serotypes (*r* = 0.8726). This means that the lower the mean pairwise identity, the higher the frequency of multi-copy, supporting multi-copy as a mechanism for gene divergence (Table 3).

We also found that the early evolution of CPS gene cluster of *V. parahaemolyticus* may be promoted by frequent multi-coping events. There are a total of 35 gene duplication events in the process of CPS evolution, whereas 26 of them (74%) are located at nodes N0 – N4 (N0:14 events, N1:6 events, N3:3 events, N4:3 events) (Table 4). Among these events, duplications of two glycoltransferase genes, *Rv2957* and *bshA*, were most active, with 8 and 7 duplication events, respectively. Notably, duplications of *Rv2957* occurred during the whole process of evolution in such hotspots as nodes N0, N3, N13, N27 (Table 4, Fig 5). In analysis on a gene-by-gene basis, the two glycoltransferase genes, *Rv2957* and *bshA*, display dual-copies at high frequencies of 15/40 (15 in 40 K-serogroups) and 9/40, respectively, and even triple copies at substantial frequencies 5/40 and 2/40, consistent with above analysis. Other multi-copies genes occur less frequently than 5 K-serogroups. In addition, other triple copies pathway gene *gtaB* and *epsM* exist at frequencies of 2/40 and 1/40, respectively (S2 Table, Fig 4, Fig 3B). Collectively, genes of multi-copies might play important roles in K antigen diversification especially through multiple pathway genes that catalyze a common precursor to different products. Notably, duplications of glycoltransferase genes, *Rv2957* and *bshA*, might play an important role in the evolution of *V. parahaemolyticus* CPS loci and are well fixed in larger amounts of K-serogroups.

We found that these mult-copies genes may have been generated by three different mechanisms in *V. parahaemolyticus*. Firstly, some genes acquire multi-copies by gene mutations. More concretely, these genes undergoes nonsense mutation leading to a stop codon, while the downstream region undergoes mutations leading to a start codon. All multi-copies of *TTHA0252* (old right-border gene) genes in five K serotypes belonging to groups 3, 4 and 5 were generated by this mechanism (Fig 4, <u>S8 Fig</u>), suggesting that these are parallel events. In addition, the gene *gtaB* in K58 has been split to two neighboring genes by nonsense mutations with respect to K6 (in group 3).

Secondly, multi-copy genes which cluster together on gene trees may arise from self-duplication. Five multi-copy genes, including 4 pathway genes (*wecB*, *wecC*, *hepC* and *kdsB*) and 1 glycoltransferase gene (*Rv2957*), are generated by self-duplication (<u>S8 Fig</u>). *wecB*, *wecC* and *hepC* duplication, with high sequence identities (72.3%, 74.1% and 41.9% respectively) uniquely occur in K17, and *kdsB* duplication with 45.1% identities, uniquely occur in K18 (Table 3), suggesting these duplication events may contribute to the origin of K17 and K18. There are 3 duplication genes in K17, which may be crucial to the development of multi-copy genes as it has the longest CPS gene cluster except K28 and K3 (Fig 2). Located on the sequence of insertion event 1 in K28 and K3, self-duplicated *Rv2957* is speculated to occur in the ancestral CPS loci of K28 and K3 and diverged along separation of K28 and K3 (<u>S8 Fig</u>). Additionally, the duplicated gene *Rv2957*, which has an opposite direction both in K28 and K3, is a border gene of insertion event 2 sequence. Thus, they may have been acquired from other species and have promote the sequence insertion by recombination.

Thirdly, apart from mutation and duplication, multi-copy genes clustered into different clades on gene trees plausibly emerge from recombination from unknown donor sources. Ten of such genes display multi-copies linked to recombination, most of which are pathway genes. Interestingly, nearly all the pathway genes with high distribution frequency achieved multi-copy by recombination, while those with low distribution tends to be generated by duplication (Table 3).

## Discussion

Our findings have provided new insights into the global evolution and diversity of CPS gene cluster of *V. parahaemolyticus*. Notably, our analysis has revealed the CPS gene cluster of 40 K-serogroups along with the finding of a conserved right-border gene, conservation in the flank region, variability in the middle region. Furthermore, we have constructed an evolution model encompassing insertion, homologous recombination, gene duplication and gene nonsense mutation to account for the observed divergence.

### Discovery of a new right-border gene of CPS gene cluster

Previous studies have suggested that CPS gene clusters of *V. parahaemolyticus* are located between *VP0214* and *VP0238* (*gmhD* – *rjg*) [27]. Our study reveals that the right-border gene *VP0238* was not conserved among K-serogroup, as revealed in this study. More specifically, 5 K-serogroups, K41, K38, K18, K28 and K3 do not contain *VP0238*. *VP0238* of another 5 K-serogroups, K20, K29, K8, K11, and K44, has undergone nonsense mutation splitting into two separated genes. In another 3 K-serogroups, K17, K19 and K32, *VP0238* is located in the middle region of CPS gene cluster (Fig 4).

We identified the gene *glpX* at downstream of *VP0238* as an accurate right-border gene of *V. parahaemolyticus* CPS gene cluster, which is conserved in all 40 K-serogroups. During our study, Yu Pang et al [42] extracted 55 K-serogroups CPS gene clusters by right-border gene *VP0238*. As we can see, for K18, K31, K38, K41, K60 which don’t have *VP0238* and K17, K19, K32 whose *VP0238* is located in the middle region of CPS gene cluster, they did not extract entire gene cluster with losing large fragment, naturally, the incorrect extraction will disturb following structure and function. What’s more, Yuansha Chen *et al.* extracted K6 CPS gene cluster losing 7 right flank genes [27]. In summary, the new right-border gene *glpX* we found in this study is helpful to build a stable method to extract entire CPS gene cluster of *V. parahaemolyticus* and benefit to following functional research and diagnosis method development.

### Evolution model of CPS loci

Study on CPS gene cluster evolution of *V. parahaemolyticus* would benefit from the discovery of a conserved right-border gene. For the first time, phylogenetic relationships of 40 K-serogroups were established in this study. Based on our evolution analysis in the results section, we propose following evolution model of CPS loci in 40 K serotypes. We speculate that all 40 K serogroups gene clusters share a common ancestor, designated as group0 for convenience, whose genetic structure is highly similar to K17. During evolutionary processes, some of group0 differentiated to group2, and others differentiated to group1 by two insertion events 1 and 2 with the duplicated gene *Rv2957*. In group1, K28 and K3 differentiated recently, as most of their genes’ position and sequence are conserved, and they have only few different genes in the middle region of gene cluster. In group 2, K17 and other two K serogroups K19, K13 differentiated by 3 gene duplication events and recombination. Subsequently, group 3 differentiated from group 2 at node 1. K6 which developed to pandemics strains are in group3. As is evident from our data, divergence of K6 and K58 was caused by mutations of the gene *gtaB*. Group 4 differentiated from group 3 at node 10 by insertion event 3, which leads to most gene cluster of group 4 and its derivative group 5 sharing three-gene sequence specially. Notably, in group4, a sequence inserted K60, K68, K36, K25 in parallel at a conserved position uniformly referred to as insertion event 5 which promoted their differentiation from others. Finally, group 5 differentiate from group 4 at node 17 by insertion event 4, and one of them K18 underwent a gene duplication event. Additionally, in general, homologous recombination occurred throughout the whole process of CPS gene cluster evolution in *V. parahaemolyticus*. This increases the diversity of CPS gene cluster structure to a large extent.

According previous reviews by Han *et al*. [21], the 24 K-serogroups, accounting for 60% of 40 in our study, were implicated in clinical pandemics of *V. parahaemolyticus* (Fig 5). These 24 pandemic K-serogroups distribution in all five groups, but were unexpectedly concentrated in group4 (Fig 5). Half of the pandemic K-serogroups belong to group 4 with high frequency at 70.5% (12/17), while frequencies for the other half are 7/13 in group 5, 3/5 in group3, 1/3 in group 2 and 1/2 in group 1. Nine K-serogroups distributed in no less than 3 countries, were recognized as wide-spread pandemic K-serogroups. Among them, 66.7% (6 in 9) are in group 4, and the other three are K3 in group 1, global spread K6 in group 3, and K12 in group 5. Our results reveal that diverse pandemic K-serogroups capable of infecting humans have polyphyletically arisen along evolution of *V. parahaemolyticus* CPS loci. The prevalence of group 4 indicates that there could be unique genetic features for synthesis specific CPS structures that enable invasion across human immune barriers. It is important to identify these genetic features and specific CPS structures to elucidate mechanisms of *V. parahaemolyticus* infection.

### Evolution mechanism

For the mechanism of variation and evolution of CPS’s structure and function, abundant literature shows that genetic variability in capsules can evolve rapidly across species by homologous recombination and horizontal transfer [33, 35–37]. Moreover, previous studies have suggested that nonsense mutations exist in polysaccharide gene clusters in some bacteria which cause one gene to split to two sections and make an enzyme nonfunctional. Examples include the c*psD* gene in CPS gene cluster of K3 serogroup in *Actinobacillus pleuropneumoniae*, and the *tyv* gene in LPS gene cluster of *Salmonella* of group A [43–46]. However, few studies have attempted to expound the effects of gene duplication in evolution of CPS gene clusters.

Elsewhere, studies have proposed that recombination might have occurred between different sister species in *Vibrio*, such as between *V. cholera* and *V. mimicus* and between *V. harveyi* and *V. campbelli* [47]. In this study, we found that not only insertion recombination, homologous recombination and gene nonsense mutation, but also gene duplication promote the evolution of CPS loci in *V. parahaemolyticus.* Other *Vibrio* species are major genetic donor sources for are the donor source inserting into CPS gene cluster (S4 Table, Fig 4, Fig 5; S6 Fig). It is worth noting that most of these donor species (*Vibrio alginolyticus*, *Vibrio campbellii*, *Vibrio harvey*) are evolutionary related, clustering in the same clade, termed the Harveyi clade by multilocus sequence analysis in a previous study [47]. *Vibrio alginolyticus*, *Vibrio campbellii*, *Vibrio harvey* have similar habitats as *V. parahaemolyticus* in seawater and some seafoods [48, 49]. Their growth on the chitinous exoskeletons of crustaceans can induce natural transformation in *Vibrionaceae* members [50]. Collectively, these suggest that recombination with neighboring *Vibrio* species living in the same ecological niches has promoted CPS gene cluster evolution in *V. parahaemolyticus*. The study and surveillance of the emergence of the novel K serotype by natural transformation between *V. parahaemolyticus* and other closed *Vibrio* species deserve attention. It can cause the serotype variation in *V. parahaemolyticus*, whereas other *Vibrio* species could increase health risks in humans by acquiring toxin-related genes from *V. parahaemolyticus*.

In our study, gene duplication is another important mechanism found promoting the evolution of CPS gene cluster in *V. parahaemolyticus*. In the early 1930s, Haldane [39] and Muller [40] were the first to propose gene duplication as a mechanism for the generation of new genes. Later, in the 1970s, Ohno substantiated the importance of gene duplication in evolution [41], and clarified how gene duplication contributes to protein functional diversity. Subsequent genomic studies have supported the notion that gene duplication is intimately associated with increased coding sequence evolutionary rates [51, 52], and regulatory sequence divergence [53]. In this study, we found that 5 multi-copy genes that have evolved through gene duplication in some or all K-serogroups with multi-copy genes. Most of them belong to pathway gene class (4/5 pathway genes, 1/5 glycoltransferase genes), distributed in K18, K17, K28, K3.

### Origin of O3:K6 serovariants

Most pandemic O3:K6 serovariants since 1996 were found to have gone through CPS loci recombination. The development of reliable methods for rapid identification of O3:K6 isolates [54] has led to the serendipitous findings of other serotypes, such as O4:K68, O1:K25, and O1:KUT (untypeable), which contain toxRS sequences, AP-PCR profiles, ribotypes, and PFGE profiles identical to those of the O3:K6 serotype [20, 28, 54]. In addition to O4:K68, O1:K25, and O1:KUT, O6:K1, which share high molecular identity with an O3:K6 isolate, were detected in Taiwan [55]. Therefore, these serotype are regarded as “serovariants” of pandemic O3:K6 [54]. Subsequently, some studies have proposed molecular mechanisms for the conversion from pandemic O3:K6 to its serovariants. Through whole-genome comparisons, Yuan Sha Chen *et al.* [56] inferred that a recombination event involving a large region of 141 kb in length covering the O-antigen and K-antigen loci occurred in pandemic O3:K6 and gave rise to the new O4:K68 serotype. However, Haihong Han *et al.* [16] postulated that the pandemic O4:K68, O1:K25, O6:K18 post-1996 originated from pandemic O3:K6 by deletion or horizontal transfer of relevant O/K antigen genes.

In contrast, based on new evidence in our study, it is more reasonable that homologous recombination of whole CPS loci between pandemic O3:K6 and non-pandemic environmental strains is a mechanism underlying the origin of pandemic O4:K68, O1:K25 and O6:K18, which has been recognized as a mode of serotype switching in *Streptococcus pneumoniae* [57] K6 and K68, K25, K18 are identified to belong to different group in our study, with K6 in group 3, K68 and K25 in group 4, K18 in group 5, and share different the genetic structure, thus the latest common ancestors of CPS loci for K68, K25, K18 are different from that of K6 (Fig 4 and 5). This means that K68, K25, K18 could not have stemmed from K6 by gene gain or loss but by CPS loci recombination between environmental K68, K25, K18 and the pandemic O3:K6 serotypes. On the other hand, our study also shows that pandemics novel K type may also arise through variations of CPS loci of pandemic O3:K6, such as pandemics O3:K58, likely has originated from O3:K6 through gene nonsense mutation. This may be another mechanism for novel serotypes/K type.

## Materials and Methods

### Bacterial culture and conventional serotyping

443 strains *V. parahaemolyticus* (S5 Table) from sentinel hospitals were collected by Shenzhen CDC. *V. parahaemolyticus* were enriched in alkaline peptone water ([pH 8.6]; 3% NaCl) and incubated at 37 °C for 16 hours on a shaker; then streaked onto *Vibrio* chromogenic agar incubated at 37 °C for 12 hours for single colonies (Guangdong Huankai Microbial Science and Technology, Guangzhou, China). Potentially productive colonies were picked and streaked onto triple sugar iron slants (Guangdong Huankai Microbial Science and Technology, Guangzhou, China) and incubated at 37 °C for 16 hours. Then subject to serotyping by serum slide agglutination tests using commercial antisera (Denka Seiken, Tokyo, Japan) according to the manufacture’s protocol and the Chinese National Food Safety Standards: Food Microbiological Examination *Vibrio parahaemolyticus* Testing, GB 4789.7-2013.

### Whole-genome sequencing and assembly

Genomic DNA were sent to BGI Research for high-throughput sequencing by using the BGISEQ-500 platform. DNA was extracted strictly according to the operation requirements of automatic nucleic acid extractor. After qualification checking using gel electrophoresis, DNA were fragmented and then processed by end repairing, A-tailing, adapter ligation, DNA size-selection, circulation, and DNA nanoball formation according to the in-house SOP of BGI library construction for BGI Seq500 (BGI-Shenzhen, China). The DNA libraries with an insert size of 300 bp were sequenced using single-end 100 bp mode (SE100) or pair-end 100 bp mode (PE100) on a BGI Seq500 Sequencer.

After quality-trimmed using SOAPnuke [58], the rest of reads were assembled into contigs by Shovill (https://github.com/tseemann/shovill) with parameter depth=0, minlen=50.

### Extraction of capsule gene cluster (CPS gene cluster)

This procedure contains coding sequence prediction, border genes identification and whole gene cluster sequence extraction. Coding sequence (CDS) of every strain was first predicted by use of prokka 1.13 [59]. Two border genes *VP0214* (gene symbol: *gmhD*) and *VP0238* (gene symbol: *rjg*) which were identified in strain RIMD 2210633 [27] are treated as capsule gene cluster border in this study. Additionally, *VP0214* and *VP0238*, adject to the above two border genes respectively, were considered as secondary border genes for more comprehensive extraction. As shown in S1 Fig, for each strain, if certain contig has one left-border gene, either *VP0214* or *VP0215*, and one right-border gene, either *VP0237* or *VP0238*, then we extract the sequence as capsule gene cluster. Border genes were queried using blast [60] with e-value less than 1e-5, identity larger than 60%, and coverage larger than 60%.

### ORFs homologous clustering, annotation and gene function classification

#### Function annotation

Opening read frames (ORFs) of CPS gene clusters function were annotated by using prokka 1.13 [59] and database selected from Swiss-Prot of UniProt. After this, if the ORFs cannot be annotated, they were blastn against the coding sequences of CPS loci of reference strain RIMD 2210633 [27] with E-value less than 1e-5, identity larger than 60%, and coverage larger than 60%.

#### Selection of representative 40 K-serogroups

To focus on different K-serogroups’ comparison, we select 40 K-serogroups’ representative strains. Firstly, compute every strains’ K antigen gene cluster length and ORF number. Afterwards, for K serogroups with more than 2 strains, the strain with K gene cluster length and ORF number equal or similar to most strains in the K serogroup was chose as the reprehensive. If K serogroup with less than 3 strains, select the representative strains at random.

#### ORFs homologous clustering and gene name assignment

To make further research in ORFs function, homologous ORFs of 40 representative strains were clustered by OrthoFinder [61] with default parameters, and assigned to orthogroups. Each orthogroups were designated as a gene groups, and were uniformly named as follows: in certain orthogroup, proportion of ORFs which were annotated using prokka, Swiss-Prot or/ reference strain RIMD 2210633, were calculated, and name of the gene with the largest proportion were chose as the name of this orthogroup. If for othogroups, all ORFs cannot been annotated as above, the orthogroup ID produced by orthofinder program were chose its name.

#### Gene function classification

To make clear their functions in capsule biosythesis pathway, genes annotated were assigned into 4 classes (according to <u>S9 Fig</u>), Pathway genes, Processing and transportation genes, Glycoltransferase genes and Function unknown or cannot been classified. The position and annotation information of each ORFs in 40 K-serogroups CPS gene clusters can be seen in S6 Table.

### Phylogenetic analysis

Species tree of 40 K were constructed based on whole length of CPS cluster using OrthoFinder with default parameters to reveal the evolution relationship among 40 K-serogroups. In addition, genes whose distribution frequency is equal to 100% in 40 K-serogroups were selected to generate core-gene evolution tree. Multiple sequence alignment were done by MUSCLE v3.8.1551 [62] with default parameter, and core gene tree were inferred using IQ-TREE [63] with parameter -m GTR+F+R9, -bb 1000, -nt 8.

### Evolution mechanism analysis

The analysis of generation mechanism of multi-copy genes are according to 3 aspects: 1.genes generated multi-copies by sequence mutation: if multi-copy genes are neighboring in a CPS gene cluster, and the sum of their length is near to the single copy gene in other CPS gene clusters, regard them as generated by sequence mutation. 2.genes generated multi-copies by gene duplication: if multi-copy genes locate at the neighboring branches in gene tree. 3.genes generated multi-copies by homologous recombination: if multi-copy genes don’t belong to 1 and 2.

The insertion events in 40 K-serogroups’ CPS gene clusters are identified by two method: Firstly, count the distribution frequency of sequence in 40 K-serogroups and observe its location feature in evolution tree. If a sequence exits in some K-serogroups which clustered in a clade of evolution tree, regard it as the potential insertion sequence. Secondly, search and identify the sequence between multi-copy genes. In addition, the source of potential insertion sequence is identified by searching on NCBI with e-value equaled to 0.

## Acknowledgements

This study was supported by National Science and Technology Major Project of China (No.2017ZX10303406), National Natural Science Foundation of China (No. 81773436), and Sanming Project of Medicine in Shenzhen (No.SZSM201811071). We thank Mingxu Li for genomic DNA preparation and serotype confirmation, and China National GeneBank for NGS library construction and sequencing. We also thank Dr. Yujun Cui for his helpful advice to this study.

## Supporting information

**S1 Fig. Capsule gene cluster extraction flow chart.**

**S2 Fig. The relationship between serotype and CPS gene cluster length, ORF number.** (A). shows the CPS gene cluster length and ORF number of every serotype grouped by K-serogroup with same color. Compared with Fig. 2, the difference is that the bars are ordered by K-serogroups. (B). shows the coding capacity and difference of CPS gene cluster of different K-serogroups. The left bar graph represents the mean length, and the right bar graph represents the mean ORF number of different K-serogroups’ CPS gene cluster. Standard deviation for which have more than one sample were indicated by error bars, and underline represents the K-serogroups whose sample number less than 3. The bars are ordered by LPS types and average length of CPS gene cluster.

**S3 Fig. CPS gene cluster of 40 K-serogroups with same gene aligned.** Compared to Fig. 4, this figure connects the same gene in neighboring K-serogroups, and only show the first occurred gene name whose function is known from top to bottom.

**S4 Fig. Presence or absence heatmap of all capsular genes of 40 K-serogroups.** Supplemental to Fig.4B, this figure shows all 217 genes of 40 K-serogroups.

**S5 Fig. Core gene tree of CPS gene cluster of 40 K-serogroups.** This evolution tree is built by core genes of CPS gene cluster. The node label is bootstrap value, and the tip label is K-serogroup.

**S6 Fig. Blast result of 5 insertion events.** This is the Graphic summary of 5 insertion events searching on the NCBI whose e-value is equal to 0. In this picture, the top 3 coverage results which are from foreign species are labeled by * with different color.

**S7 Fig. The gene trees of insertion event 5.** This is the gene trees of seven genes in insertion event 5. In every tree, tip labels refer to K-serogroup and gene id.

**S8 Fig. 16 gene trees that contain multi-copies.** This is the gene trees of 16 genes which contain mult-copies. In every tree, tip labels refer to K-serogroup and gene id. What’s more, the red font with red box indicate the multi-copies originated by gene-self duplication, the blue and orange font refer to multi-copies originated by gene nonsense mutation and recombination apparently.

**S9 Fig. Gene function classification flow chart.**

**S1 Table. Length and ORF number status of CPS gene cluster in 40 K-serogroups.**

**S2 Table. Classification, identity and distribution frequency of 40 K-serogroups genes.**

**S3 Table. The location of duplication events on evolution tree.** Every row shows a duplication event of gene. The column with * shows the node location of duplication event on evolution tree (as figure 5 shown).

**S4 Table. Blast results of insertion events.** Note: Sheet 6 summarizes the top 3 blast results ordered by score from other *Vibrio* species of 5 insertion events.

**S5 Table. Strain information in this study.** Note: The strain, in which entire CPS gene cluster could be extracted, are indicated with *, namely the flank genes *VP0214* and *glpX* exits on a same contig.

**S6 Table. ORFs information of CPS gene clusters in 40 K-serogroups.**

### Data reporting

The data that support the findings of this study have been deposited in the CNSA (https://db.cngb.org/cnsa/) of CNGBdb with accession code CNP0000343. The assembly accession numbers are listed in Table S5.

